# Where the present gets remembered: Sensory regions communicate with the brain over the longest timescales

**DOI:** 10.1101/2023.09.18.558347

**Authors:** Greg Cooper, George Blackburne, Tessa Dekker, Ravi K Das, Jeremy I Skipper

## Abstract

In natural contexts, the brain simultaneously processes sensory information over diverse timescales. Here we reveal how this is reflected in the organisation of asynchronous connectivity in the brain. In 86 individuals watching feature-length movies during functional neuroimaging, we calculated the delay at peak connectivity between brain regions. We found the longest delays in received whole-brain functional connectivity within ‘sensory’ regions (:S 18 seconds). Two complementary dimensionality reduction approaches were used to probe the spatial organisation of connection delays and weights. First, clustering of received delays separated sensory, and transmodal/associative outputting regions, suggesting that putatively localised functions are associated with asynchronous local-to-whole-brain connectivity patterns. Next, we organised delayed connectivity maps by likeness, unveiling five orthogonal gradients of variation, each demonstrating associations between early-sensory and transmodal/associative regions. Together, these findings challenge contemporary conceptualisations of the brain’s temporal hierarchy by emphasising the role of sensory regions as sites of integration across timescales.

## Introduction

The brain must extract information from sensory input across various timescales, ranging from millisecond-level changes (e.g., in visual processing) to gradual shifts (e.g., in topics of conversation)^1–3^. This process is not well understood. On the one hand, the brain appears to be able to track fast and slow streams of information independently. For instance, it is possible to attend to the individual words in a given sequence, without losing track of the meaning of the sentences that they form. On the other hand, the brain must also be able to leverage slowly evolving contextual information to constrain the processing of ambiguous sensory inputs presented over shorter timescales^4–7^. For example, the meaning of ‘bat’ depends on whether the conversation is about animals or cricket.

Consequently, the real-time processing of multiple timescales of information at once entails mutual dependencies between processing timescales; whereby shorter units of information combine to form longer units, but not before the shorter units have been shaped by longer prior units. How then does the brain maintain appropriate sensitivity to multiple different timescales of sensory information, while allowing for mutually constraining interactions between them?

Contemporary views on cognition across timescales largely support a process through which information is constructed in sequential steps of increasing duration across a hierarchy of cortical regions^1,8,910^. In this model, ‘unimodal’ early sensory regions (e.g., primary visual and auditory cortices) preferentially process the shortest timescales of information, while longer timescales are processed in ‘transmodal’ regions, such as the so-called default mode network^11^, associated with higher-order cognitive functions such as autobiographical memory. This progression of timescales up the hierarchy appears to persist in the absence of stimulation^12,13,14^, and under various task states^15^, suggesting that early sensory regions intrinsically operate at faster timescales than higher-order regions^3,16,17^.

Coordinated activity between distant brain regions likely plays a key role in shaping their preferred timescales of processing. For instance, in a computational model of the macaque cortex, Chaudhuri et al demonstrated that disrupting long-range connections attenuates local response timescales and disrupts the hierarchy across regions^18^. Further, the anatomical distribution of timescales closely aligns with the principal gradient of cortical connectivity, which similarly runs from unimodal to transmodal regions^19–21^. Thus, the lengthening of processing timescales along a uni-to-transmodal gradient of connectivity implies the perspective that transmodal regions are uniquely capable of integrating contextual information from temporally diverse sources^20^ because of their dissimilarity to the whole-brain connectivities of more rapidly fluctuating sensory regions.

Interpreting these apparent hierarchical distinctions between unimodal and transmodal regions within a predictive-processing framework helps to shed light on why early sensory processing regions are particularly sensitive to fine-grained, moment-to-moment information^22^. Because these neocortical regions are the closest to peripheral sensory organs in terms of synaptic-distance, they are well placed to compare and update their current ‘predictions’ to the temporally fine-grained sensory input they receive. However, these predictions do not form in isolation. Rather, they are shaped by feedback from higher-order brain regions, which are able to incorporate information from across a range of timescales^11,16^. This feedback constitutes the ‘priors’ or anticipatory models generated by the brain, based on previously stored and processed information. Hence, feedback plays a critical role in shaping the functional specialisation of sensory regions.

Indeed, the role of feedback processing in auditory and visual contexts is well evidenced. Spoken language often contains ambiguity in its ‘noisy’ acoustic features, words, and sentences. Consequently, real-world language comprehension relies on contextual information encoded over longer timescales to interpret ambiguous speech in the auditory cortex^4–7^. In the visual domain, resolving the ambiguity of retinal inputs does not depend solely on detection of low-level stimulus features^23^, but employs extensive top-down processing constraints, as reviewed by Petro *et al* ^24^. This is reflected in the fact that −70% of the auditory system’s connections stem from other cortical regions^25,26^ which is mirrored by the preponderance of incoming, over outgoing, connections into early visual processing regions ^27–31^.

This feedback-dominated model, wherein the specificity of early sensory regions to their preferred sensory modality is contingent on the feedback of prior information pertaining to multiple timescales of information appears to contradict the feed-forward picture of cognition across timescales^20,32^, wherein information is aggregated over increasing temporal windows along the uni-to-transmodal cortical gradient. This might, however, be reconciled by considering that these two processes are complementary. Sensory information does, indeed, feed-forward along a cortical gradient, but this is not a one-way process. Instead, it is shaped and refined by continuous top-down influences from higher-order brain regions.

We therefore hypothesize that ‘early’ sensory regions sit at both the beginning and the end of a hierarchy of temporally extended cortical processing. Consistent with feed-forward accounts of hierarchical temporal processing, we expect motor, somatosensory, primary visual, and primary auditory regions to transmit information to the rest of the brain over the longest timescales, reflecting the accumulation of contextual information constructed from temporally fine-grained information outwards from these regions. We also expect these regions to *receive* information from the rest of the cortex over the longest and most diverse range of timescales, satisfying their presumed requirement for feedback from various sources specified by predictive processing models of brain function.

We test this hypothesis in a group of participants undergoing blood oxygenation level dependent (BOLD) functional magnetic resonance imaging (fMRI). with feature-length movie stimuli that emulate everyday ecological temporal processing demands, where information evolves over different timescales ^33–36^. To do so, we investigate the delayed connectivity architecture across brain regions by employing a cross-correlation approach that enables rapid calculation of delayed communication between regions over long timescales in the order of tens of seconds, at sub-repetition time (TR) resolution from fMRI data (Figure 1)^37^. In practice, we generate estimates of the delay at which cross-correlation is maximal between a given brain region and the rest of the cortex (Figure 1A, 1B), and subsequently aggregate these ‘delay maps’ to create new indices of regional ‘input delay’ and ‘output delay’ (Figure 1C).

**Figure 1.**
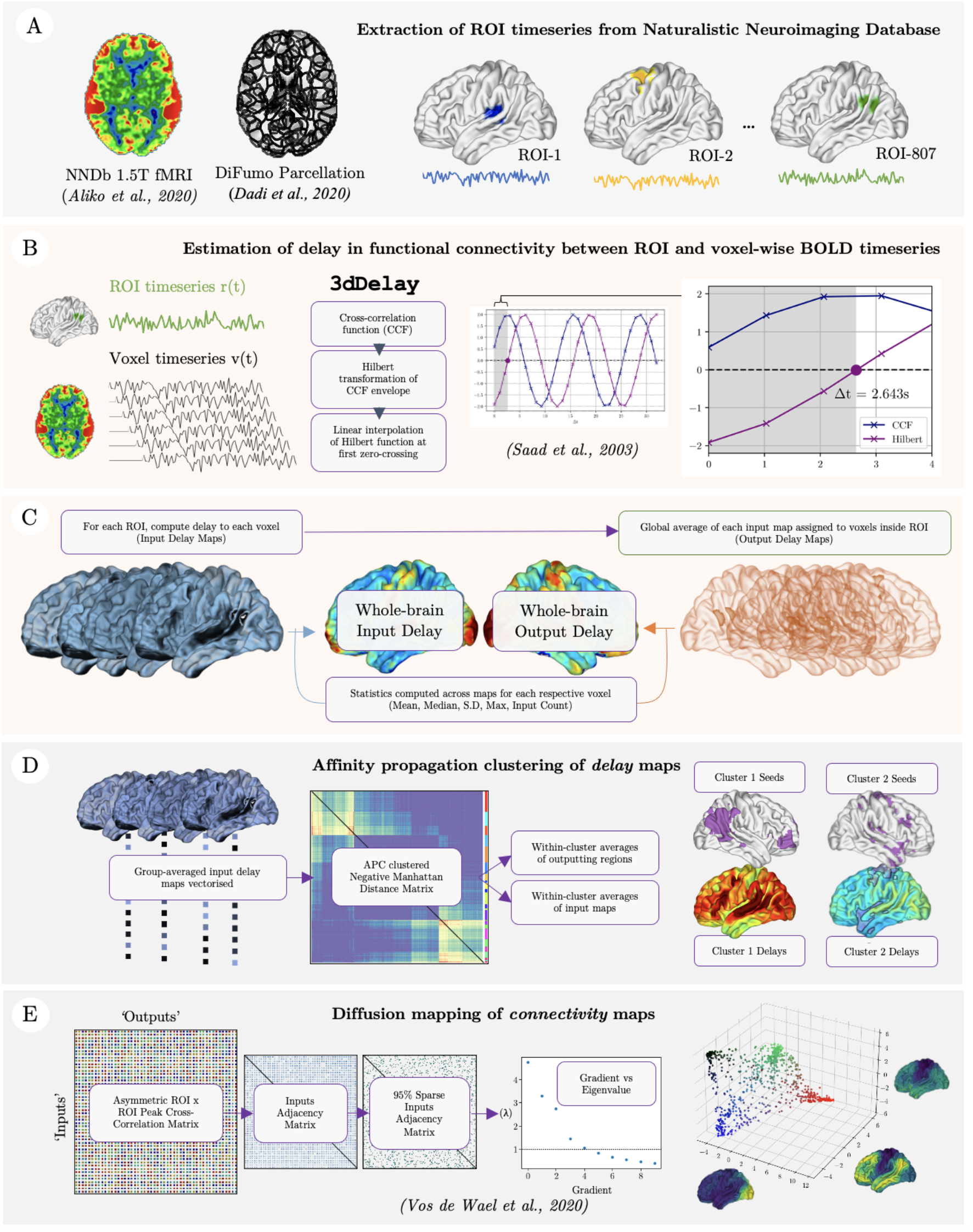
Delay mapping. A) ROT parcellation and signal extraction from each of 86 participants, in three separate 20 minute chunks of movie time course, representing the beginning, middle and final sections of each movie in order to assess the test-retest reliability of our measure within participants. B) Region of interest (ROT)-wise estimations of the delay at which the cross correlation coefficient between extracted ROT signals and voxel time series is highest in each voxel, yielding one ‘delay-map’ per ROT, per participant, per chunk. To remove spurious or low-strength delayed connections, these maps are then thresholded, as detailed below. C). Each ROT-wise delay map is concatenated for subsequent, participant-level, aggregate statistical analyses, which are computed across ROT-wise maps (’Whole-brain Tnput Delay). To estimate the average delay over which each given ROT is transmitting information to the rest of the cortex (’Whole-Brain Output-Delay’) are calculated for each ROT, and aggregated to yield a map of the average delayed global output of each region. D). Designation of ‘networks’ of delayed connectivity is achieved via affinity propagation clustering of vectorised group-averaged delay maps. E). The characterisation of ‘Gradients’ of delayed connectivity are computed via diffusion embedding of similarities between ROT-wise peak cross-correlation coefficients.

As we anticipate that each ‘outputting’ seed brain region will manifest a distinctive spatial distribution of connection delays, we hypothesise that networks tied to different cognitive functions - such as audition, language processing, visual processing, and other processes not linked to any one sensory modality - will display spatially separable patterns of delayed global output connectivities. These patterns may offer insights into the temporal encoding requirements of each of these functions. We therefore use a data-driven technique to decompose maps of the delay in seconds over which each region receives input from an outputting seed region, into a set of aggregated networks of outputting seed regions. We expect this to unveil an underlying organisation that reflects a variety of temporal connectivity patterns within and across sensory modalities, canonical higher-order networks, and subcortical regions (Figure 1D).

Gradient mapping encompasses a complementary dimensionality-reduction technique to understand the organisation of connections across the brain^38^. Unlike clustering techniques, gradient mapping can reveal multiple continuous gradients of variation in the whole-brain functional connectivity of each region by ordering the connectivity profile of each region along orthogonal dimensions according to the geometric distances between their spatial distributions. To resolve the hypothesised heterarchical (as opposed to hierarchical) organisation of sensory systems in the brain^39–41^ we extend the application of gradient mapping of functional connectomes^38^ acquired during naturalistic viewing^21^ to reveal where delay-adjusted whole-brain connectivity patterns are similar between regions (Figure 1E). In the context of whole-brain interactions under naturalistic conditions, we expected to observe close associations between sensory and ‘higher-order’ regions, reflecting their respective roles as sites of integration of temporally extended processes.

## Results

### Whole brain delayed connectivity

Our primary hypothesis posits that primary sensory regions receive brain-wide inputs over extended timescales under naturalistic conditions. To test this, we compute the delay-at-peak cross-correlation coefficient between the time series of each of 807 Regions of interest (ROIs) covering the brain, with every other voxel in the brain, generating 807 ROI-wise ‘input-delay’ maps (Figure 1B). To summarise the intraregional delayed input connectivity of each voxel in the brain, we then compute the voxel-wise statistics over every delay map (Figure 1C), including non-zero means, medians, standard deviations and 99th percentile of delay values (to provide a conservative estimate of the longest delays over which communication occurs between regions), as well as the number of voxels that survive thresholding (R > 0.1) across all ROI-wise input-delay maps (to estimate the node-centrality of a region over delayed connections).

Applying this to a dataset of 86 individuals viewing one of 10 movies, we examine the group-level statistics describing the delay at which each region of the cortex receives inputs from, and transmits information to the rest of the brain. To account for contextual variation in naturalistic stimuli, we examine this over three 20 minute segments covering the beginning, middle and end of movies. To do so, we employ a linear mixed effects model considering gender, age, watched movies, and segments.

Figure 2 displays the input connectivity-delay maps collapsed over segments, revealing patterns of delayed communication in response to dynamic stimuli that are largely in-line with our predictions. We observe relatively low median delay in the basal ganglia and insula (Figure 2.A.i), while the highest medians, means, standard deviations and 99th percentile of delays, are present in the central sulci (encompassing the primary motor and somatosensory cortices), calcarine sulci (commonly referred to as the primary visual cortex), bilateral transverse temporal gyri (primary auditory cortex), and neighbouring superior temporal regions, including the plana polare and temporale, and superior temporal gyri.

**Figure 2.**
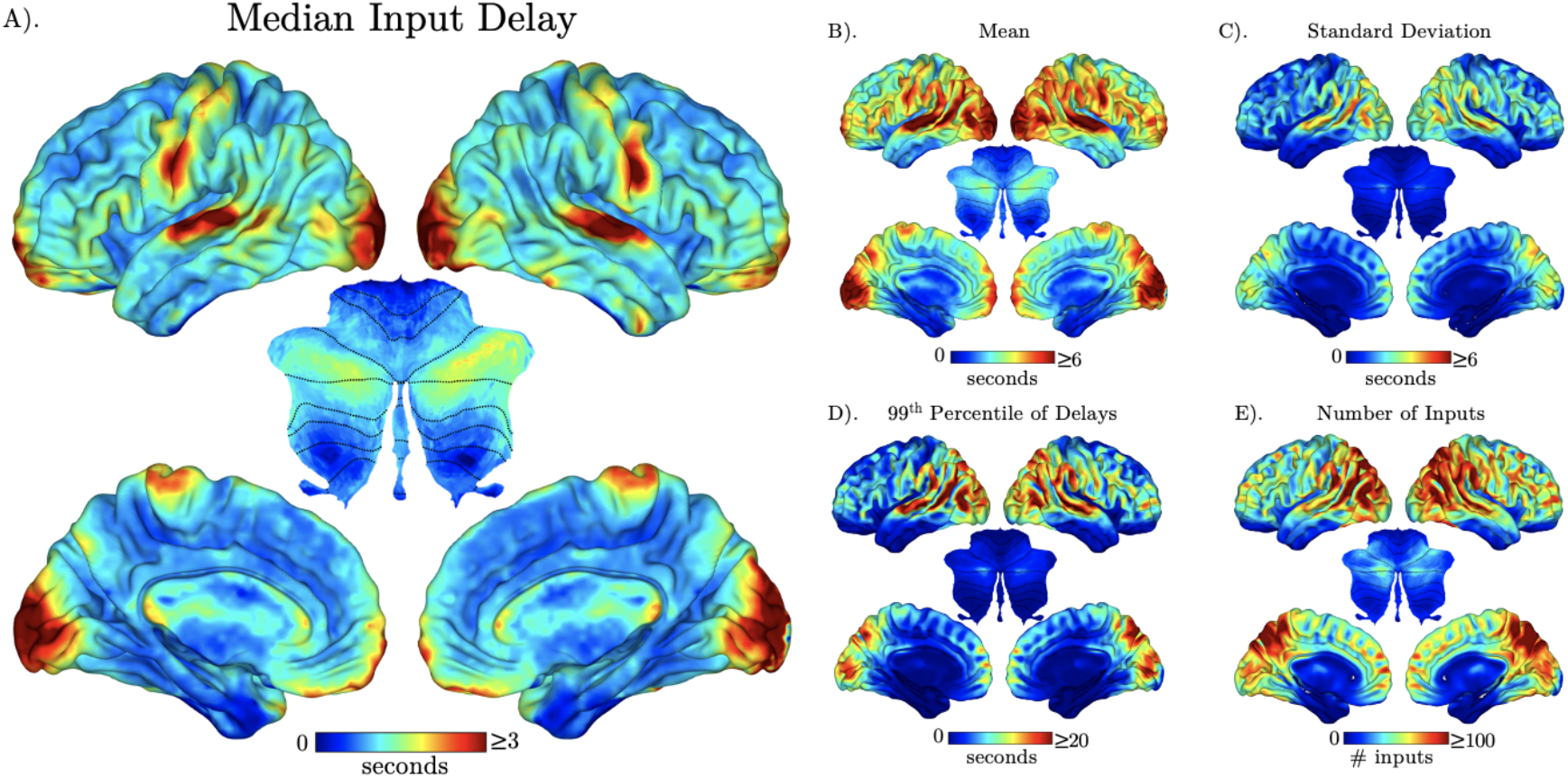
Delayed connectivity maps. A group-level linear mixed effects model was used to generate non-zero median input delay maps in seconds across three movie segments (A), mean (B), standard deviation (C), 99th percentile (D), and the number of inputs received by a voxel from other ROTs where the cross-correlation coefficient R ≥ 0.1 (E). All results are thresholded, corrected for multiple comparisons at alpha = 0.01.

Similarly, output maps were generated by computing the global statistics of each ROI-wise delay map, and projecting these values back into the voxels of each respective ‘outputting’ ROI. These maps demonstrated similarly high, mean and median delays in primary sensory regions, but with a higher standard deviation (Table 1) and a more diffuse distribution around these areas (Supplementary Figure S1; see Supplementary Figure S2 for analyses of individual movie segments).

**Table 1.** High-delay clusters. Cluster data generated by thresholding median delayed input maps at the 99th percentile of observed delay values (≥ 2.30 seconds). Estimates of the voxel-wise global output delays across the whole brain are presented in supplementary Figure S1.

| Cluster | 1 | 2 | 3 | 4 | 5 | 6 | 7 |
| --- | --- | --- | --- | --- | --- | --- | --- |
| Voxels | 1980 | 383 | 231 | 230 | 182 | 126 | 125 |
| Peak XYZ (RAI) | -4.5 94.5 28.5 | 25.5 -67.5 -10.5 | 61.5 16.5 7.5 | -64.5 10.5 4.5 | -1.5 -40.5 55.5 | 55.5 7.5 31.5 | -58.5 1.5 28.5 |
| Region label (glasser) | R_Primary_Visual_Cortex | R_Area_10v | L_Auditory_5_Complex | R_Auditory_5_Complex | R_Supplementary_And_Cingulate_Eye | L_Primary_Sensory_Cortex | R_Primary_Motor_Cortex |
| Author label | Primary Visual Cortex | Middle Orbital Gyrus | Left Superior Temporal Gyrus | Right Superior Temporal Gyrus | Left Supplementary Motor Area | Left Postcentral Gyrus | Right Postcentral Gyrus |
| median IN (secs) | 2.99505 | 2.65188 | 2.8656 | 2.88817 | 3.13566 | 2.7767 | 2.78065 |
| median OUT (secs) | 2.88777 | 1.75504 | 2.13169 | 2.25828 | 1.57529 | 2.04902 | 2.35677 |
| mean IN (secs) | 5.62354 | 4.4807 | 7.12086 | 6.91959 | 4.92535 | 5.60241 | 5.46165 |
| mean OUT (secs) | 5.41319 | 4.50279 | 5.13252 | 5.30406 | 3.54891 | 4.52308 | 4.57286 |
| stdev IN (secs) | 2.2224 | 1.27357 | 3.84382 | 3.5278 | 1.87311 | 2.53548 | 2.37977 |
| stdev OUT (secs) | 7.55355 | 8.07687 | 8.43502 | 8.46312 | 5.93647 | 6.83011 | 6.34765 |
| 99th percentile IN (secs) | 9.57582 | 3.67788 | 17.9958 | 16.0443 | 7.03918 | 11.4391 | 10.7189 |
| 99th percentile OUT (secs) | 31.4107 | 34.783 | 36.2956 | 37.0254 | 25.3632 | 29.1509 | 26.9265 |
| count IN (number of inputs above threshold) | 47.3616 | 24.2066 | 70.1455 | 63.6776 | 52.6981 | 56.8533 | 54.3738 |

### Recovery of Large-Scale Delay Networks

The voxelwise statistical aggregation (e.g., median delay) of every delay map originating from separate regions across the brain obscures separable spatial patterns of connectivity delays that contribute to the observed global patterns. Given that ecological sensory information comprises multiple separable sets of sense-specific timescales, we expect that regions that are associated with separable cognitive functions to exhibit meaningfully separable clusters of delayed global *output* connectivities (i.e., spatially similar output-delay topographies) that are reflective of their shared temporal encoding demands. Specifically, we expect delay maps originating from regions associated with audition and language, vision and attention, and somatosensation to form separable clusters.

To test this in a data-driven manner, while allowing for groupings of outputting seed-regions of variable sizes in reflection of the heterogeneity of sensory processing systems, we employed Affinity Propagation Clustering (APC), a dimensionality reduction technique, which unlike k-means doesn’t require the arbitrary a priori specification of a number of clusters. We applied APC to ROI-wise delay maps averaged over participants and movie chunks, cutting the resulting dendrogram at the median of input preferences^42^ to capture a range of clusters comprising varying numbers of seed ROIs (Figure 1D). The ROIs from which each cluster was generated were then recovered and superimposed onto cluster-wise delay maps to generate estimates of the functional network giving rise to the delay topography of each cluster.

This revealed 24 distinct sets of regions that generate separable topographies of delayed connectivity. Manual inspection of these clusters, assisted by performing a topic-wise meta-analysis of these regions against the NeuroSynth database^43^ facilitated their manual segregation into 5 anatomical domains, representing networks associated with 1) audition and language, 2) vision and attention, 3) motor processing and somatosensation, 4) associative and higher-order processes, and 5) subcortical regions (Figure 3A). Neurosynth topic-wise decoding was applied to maps of the outputting ROIs that generated the delay topography of each presented cluster and supports this subjective grouping of clusters (Figure 3B).

**Figure 3.**
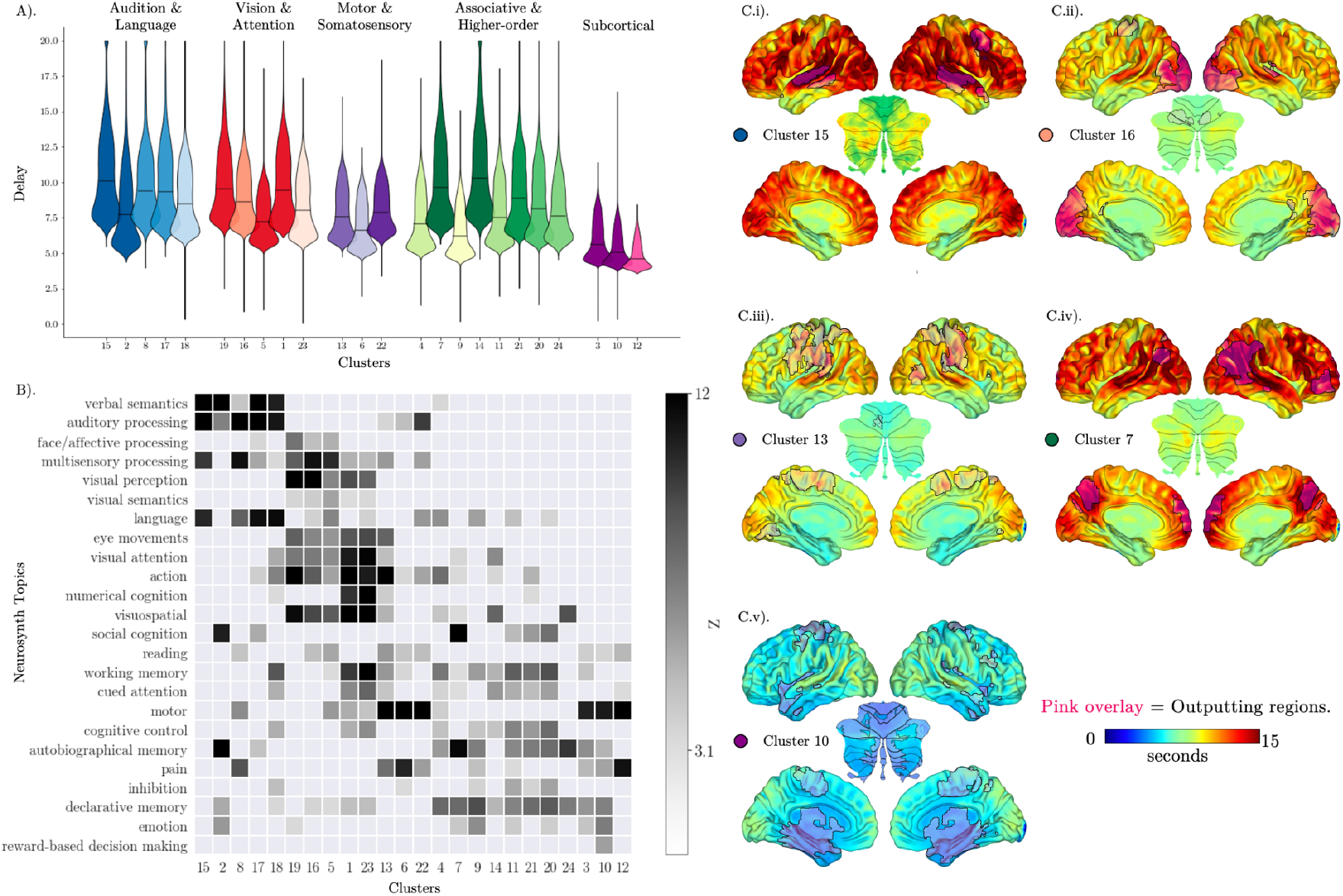
Clustering of ROT-wise delayed connectivity maps by Affinity Propagation Clustering. A). Distribution of voxel-wise delay values in each cluster-map (labelled as clusters 1-24, along the X-axis). Each of the 24 clusters are manually grouped into one of five domains (labelled at the top of the violin plot as ‘Audition and Language’, ‘Vision & Attention’, ‘Motor & Somatosensory’, ‘Associative & Higher-order’ and ‘Subcortical), by interpretation of the associated cognitive functions and anatomical distribution of the outputting ‘seed’ regions responsible for producing the connectivity-delay topographies comprised by each cluster. This manual categorisation process is assisted by a meta-analysis showing the association between a list of 24 topic terms derived from the Neurosynth database, with binary masks of the outputting seed regions of each cluster (B). Examples of clusters from each of the five domains are depicted in C.i-v), where the recovered outputting seed regions are superimposed in pink over the clusters of delayed connectivity maps that they generate. Each cluster (B) described in detail in table 2. Plots of the remaining 19 clusters, their decoding terms, and relevant statistics are available in Supplementary Figures S4, S5 and Supplementary Table T2.

The distribution of global delayed connectivity values varied between clusters within each category. However, the highest global median output delays were observed within clusters originating from ‘audition and language’, and ‘associative and higher-order’ networks (Figure 3A, Ci). The lowest values were consistently observed within clusters originating from subcortical regions (Figure 3A, C.v; Table 2). Plots of the remaining 19 clusters, their decoding terms, and relevant statistics are available in Supplementary Figures S4, S5 and Supplementary Table T2.

**Table 2.** Summary of delayed connectivity networks yielded by Affinity Propagation Clustering. Plots of the remaining 19 clusters, their decoding terms, and relevant statistics are available in Supplementary Figures S4, S5 and Supplementary Table T2.

| **Example Cluster & manually-assigned domain** | **Cluster 15**<br>**(Audition & Language)** | **Cluster 16**<br>**(Vision & Attention)** | **Cluster 13**<br>**(Motor & Somatosensation)** | **Cluster 7**<br>**(Associative & Higher-order)** | **Cluster 10**<br>**(‘Subcortical’)** |
| --- | --- | --- | --- | --- | --- |
| **Outputting Brain Regions** | STG, STS, right middle frontal gyrus | Occipital cortex, right Heschl’s gyrus, anterior middle portion of the postcentral gyrus | Pre and post-central gyri and sulci, left parietal operculum, left STG, left lingual gyrus | Precuneus, angular gyri and sulci, medial frontal pole, orbital cortex | Basal ganglia, cerebellar lobule VI and Crus I, amygdalae, hippocampi, temporal poles, inferior portions of insula, occipitotemporal gyrus, medial portions of precentral gyri and sulci |
| **Global Median Delay ± Standard Deviation** | 10.1 seconds ($\pm 2.89$ seconds) | 8.94 seconds ($\pm 1.82$ seconds) | 7.59 seconds ($\pm 1.68$ seconds) | 8.89 seconds ($\pm 2.29$ seconds) | 4.58 seconds ($\pm 0.89$ seconds) |
| **Neurosynth Topics** | 'verbal semantics', 'auditory processing', 'multisensory processing', 'language' | 'face/affective processing', 'multisensory processing', 'visual perception', 'visual semantics', 'eye movements', 'visual attention', 'action', 'visuospatial', 'declarative memory' | 'action', 'motor', 'pain', 'multisensory processing' | 'social cognition', 'autobiographical memory', 'declarative memory', 'emotion' | 'motor', 'autobiographical memory', 'pain', 'declarative memory', 'emotion', 'reward based decision making' |
| **$\leq 5$th Percentile Regions and Delay** | Bilateral subcortical regions including putamen, thalamus, hippocampus, amygdala, cerebellar lobule V (2830 voxels, mean delay = 6.89 seconds) | Subcortical cluster including basal ganglia, hippocampi, amygdala, cerebellar lobule V, middle cingulate cortex (3277 voxels, mean delay = 6.46 seconds) | Bilateral subcortical structures including hippocampus, parahippocampal gyri, thalamus, pallidum, caudate nuclei, cerebellar lobules IV & V (3325 voxels, 5.71 seconds) | Basal ganglia (3390 voxels, 6.66 seconds) | Basal ganglia (3475 voxels, 4.04 seconds) |
| **$\geq 95$th Percentile Regions (voxels, mean delay)** | Right STG (1117 voxels, 18.6s), left STG (1099 voxels, 25.53s), right inferior frontal gyrus extending to right precentral gyrus (482 voxels, 20.94s), left inferior frontal gyrus (194 voxels, 16.41s), left precentral gyrus (143 voxels, 17.72s), right superior occipital gyrus (117 voxels, 16.01s), left middle occipital gyrus (77 voxels, 17.27s) | Calcarine gyrus, right calcarine, bilateral middle occipital gyri, left STG and MTG (2905 voxels, 12.85s), right STG (249 voxels, 12.691s) | Bilateral STG, MTG, calcarine, middle and superior occipital gyri, cuneus (2859 voxels, 11.35s), frontal superior medial gyrus | Right superior temporal sulcus, MTG, angular gyrus, temporoparietooccipital junction (1142 voxels, 16.24s), right middle, medial, and inferior frontal gyri (1035 voxels, 15.8s), left middle temporal and angular gyri (876 voxels, 16.24s), precuneus (208 voxels, 15.55s) | Bilateral STG, MTG, angular gyri, middle and superior occipital gyri, calcarine gyri, precuneus (3006 voxels, 7.92s), superior medial gyri (126 voxels, 7.71s), right precentral gyrus (105 voxels, 7.81s) |

### Delayed Functional Connectivity Gradients

Under resting state conditions, the principal gradient of connectivity places primary sensory regions at a maximal distance from the default mode network ^20^, where connectivity is defined as instantaneous correlation between ROI timeseries. Under naturalistic stimulation, cortical gradients can be separated by sensory modality ^21^, though higher order processing networks remain uniformly at a maximum distance from sensory regions ^20,21,38,44,45^. We sought to reassess the principal gradients of connectivity under more ecologically valid contexts (using naturalistic stimulation) and allowing for delays in directed connectivity between regions, by computing the peak correlation between regions across delays. We do so to test the hypothesis that accounting for delays in direct connectivity between brain regions results in a close association between sensory and higher-order regions along connectivity gradients, reflecting their respective roles as sites of integration of temporally extended information.

ROI diffusion embeddings were determined using delay-adjusted input connectivities, approximated as a delayed Pearson’s R. Only the first five incoming-delayed-connectivity gradients with eigenvalues > 1 were considered significant (see supplementary Figure S5). Figure 4A displays a 3D scatter plot, where each point represents a region of interest and its connection patterns with other ROIs. The colour of these points reflects their connectivity characteristics, allowing a visual representation of connectomic similarities. Within this 3D space, ROI embedding coordinates along the first three gradients are arranged within an approximately tetrahedral structure, as made apparent by the opposing coloration of vertices in red (denoting ROIs largely centred around the occipital cortex), black (featuring midbrain ROIs covering the basal ganglia), green (frontal and lateral temporal lobes) and blue (ROIs centred around the central sulcus). To provide functional attributions to the regions in figures along each gradient, a neurosynth topic-level meta-analysis of binary masks taken from each 5-percentile-band of each gradient are presented alongside surface renderings (Figures 4.C-G.i).

**Figure 4.**
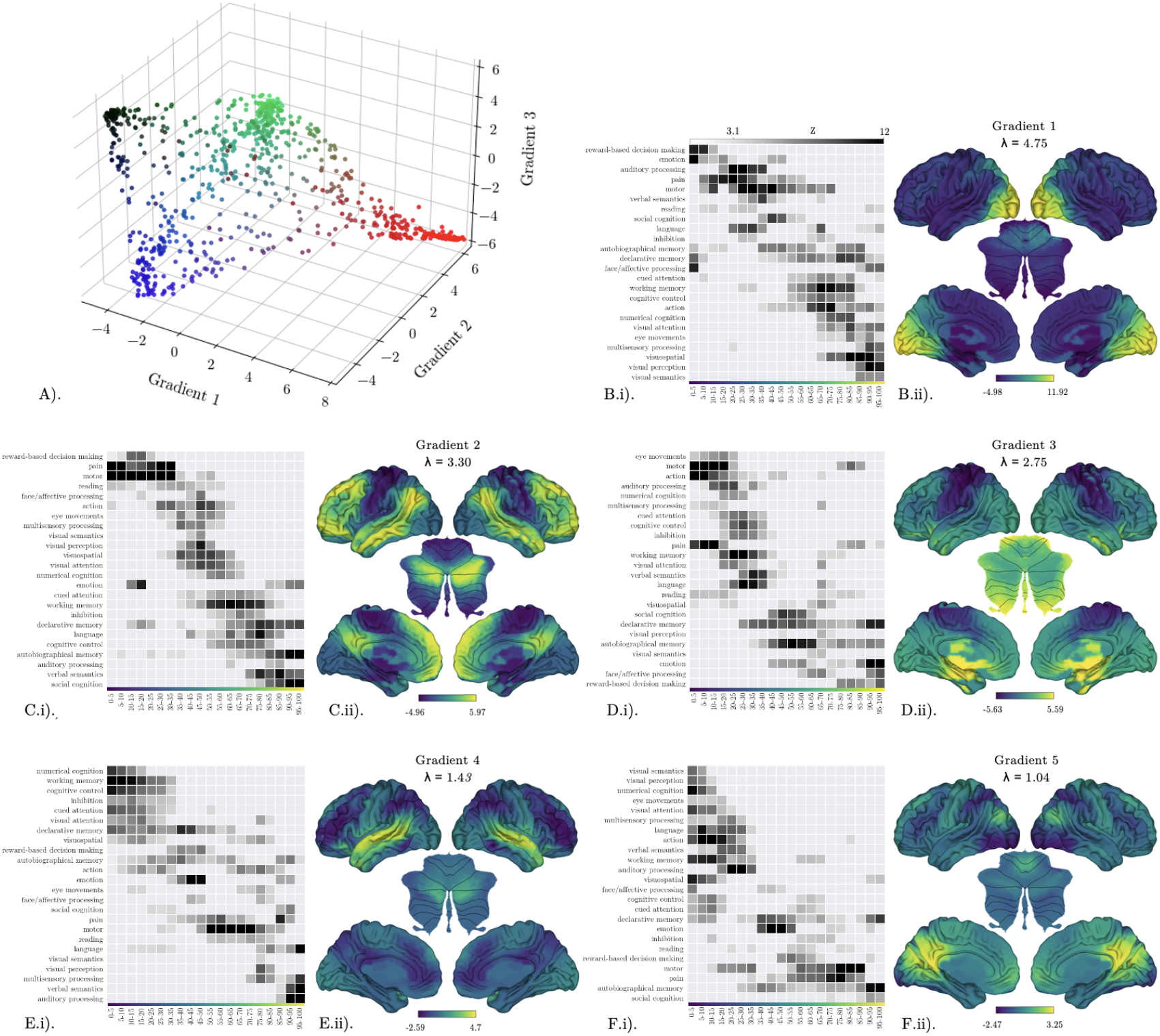
Diffusion mapping. 3D embedding of the input connectivities received by each ROT, where each ROT has been assigned an RGB value relative to their position along the first three diffusion gradients. B-F.i.) Neurosynth meta-analytic topic-wise decoding matrix of 5-percentile bins along gradients 1-5, respectively. B-F.ii). Surface plots of gradient 1-5, respectively, where the embedding coordinate value of each ROT is pro}ected back onto the cortical surface, arranged in descending order of eigenvalue (J).

Unique combinations of closely embedded sensory and transmodal regions can be observed in each gradient. For example, in gradient one (Figure 4B), the yellow pole comprises the calcarine sulci and the occipital cortex more generally, and is associated with meta-analytic topics such as ‘visuospatial’ and ‘visual perception’. These regions extend to areas like the putamen, posterior cingulate cortex and angular gyri, linked to diverse higher-order functions like ‘autobiographical memory’ and ‘visual attention’. Meanwhile, the blue pole includes the basal ganglia, superior temporal gyri, and central sulci and its functions range from ‘reward-based decision making’ and ‘emotion’ to ‘auditory’ and ‘motor’. This trend of sensory-to-higher-order pairings is continued across gradients. For instance, gradient two (Figure 4C), which at one extreme depicts similar embeddings between regions associated with auditory and language processing, such as the superior temporal gyrus and inferior frontal gyrus, with canonical nodes of the default mode network including the posterior cingulate, angular, and medial temporal gyri. In gradient three (Figure 4D, language-associated and visual processing associated nodes are positioned closely to transmodal nodes in the centre of a motor-to-basal-ganglia axis. Gradient four (Figure 4E) reveals an axis of higher-order-cognition to audition/language/vision, anchored on one end by a group of regions resembling the frontoparietal attention network^46^, and associated with the functions of ‘numerical cognition’ and ‘cognitive control’, and on the other end, in the superior temporal gyri and occipital cortex, being associated with the topics of ‘auditory processing’, ‘verbal semantics’, ‘multisensory processing’ and ‘visual perception’. This gradient was centred around visual, motor, and subcortical/limbic (’emotion’, ‘reward-based decision making’) topics. Finally, gradient five (Figure 4F) reveals close embeddings between regions associated with the topics of ‘autobiographical memory’, ‘social cognition’, ‘motor processing’ and pain. Complete descriptions of remaining gradients are provided in Table 3.

**Table 3.** Summary of delayed input connectivity gradients, including the eigenvalue of each gradient (A) a description of the regions along the gradient falling within the 70th-100th percentiles, the 30th to 70th percentiles, and the 0th to 30th percentiles, as well as a list of decoded meta-analytic topics.

|  | *Gradient 1* | *Gradient 2* | *Gradient 3* | *Gradient 4* | *Gradient 5* |
| --- | --- | --- | --- | --- | --- |
| Gradient Eigenvalue ($\lambda$) | 4.75 | 3.30 | 2.75 | 1.43 | 1.04 |
| Anchored Regions (70th-100th percentiles) | Calcarine gyri, Occipital cortex, Fusiform gyri, Precuneus, Posterior cingulate cortex, angular gyri | Cerebellar Crus I & II, Precuneus, Posterior cingulate cortex, Orbital/Medial/Inferior frontal cortices, Angular/Inferior/Middle temporal gyri, Superior temporal sulcus | Basal ganglia, Cerebellum, Medial/Inferior temporal regions, Inferior temporal gyri, Hippocampus, Fusiform gyri | Temporal cortex, Left inferior frontal gyrus, Temporal poles, Anterior inferior/middle temporal gyri, Superior temporal sulci/gyri, Temporoparieto-occipital junction, Cerebellar crus I & II | Precuneus, Posterior cingulate, Medial prefrontal cortices |
| Equidistant Regions | Angular gyri, Superior parietal lobules, Cerebellar crus I & II | STG, Heschl's gyrus, Planum temporale, Early visual areas, Occipital cortex, Middle & anterior cingulate cortex, Anterior insula, Pars opercularis | Frontal, Parietal, Occipital cortices, STG, MTG | Occipital cortex, Pre and post-central gyri, Orbitofrontal cortex, Medial temporal gyri, Hippocampus, Parahippocampal/Fusiform gyri, Basal ganglia | Superior frontal, Pre and post-central gyri, Rolandic operculae, Middle/Superior temporal cortices, Insulae, Anterior cingulate cortex, Basal ganglia, Entire cerebellum (except Crus II) |
| Opposite Pole (0-30th percentiles) | Subcortical structures (Hippocampi, Parahippocampal gyri, Putamen, Caudate, Striatum, Thalamus), Cerebellar lobules IX and X, Insular cortex, Frontal and Temporal lobes | Cerebellar lobules V & VIII, Basal ganglia, Medial temporal structures, Pre and post-central gyri | Pre and post-central gyri | Anterior/Posterior cingulate cortices, Inferior/Superior parietal lobules, Inferior/Middle frontal gyri | Early visual cortex, Calcarine/Lingual gyri, Fusiform gyri, Posterior inferior temporal gyri, Superior/Inferior parietal lobules |
| Functional Decoding (Percentile Bands) | 0-30: Reward, Emotion; 30-70: Auditory, Sensorimotor, Pain; 70-100: Language, Autobiographical memory, Visual perception | 0-30: Somatosensation, Motor control; 30-70: Visual processing; 70-100: Language, Cognitive control, Autobiographical memory, Auditory processing, Verbal semantics, Social cognition | 0-30: Motor, Somatosensory, Auditory; 30-70: Higher-order cognitive functions, Visual processing; 70-100: Affect, Reward-based decision making | 0-15: Cognitive control, Inhibition; 15-85: Reward, Social cognition, Autobiographical memory, Motor, Somatosensory, Pain, Visual; 85-100: Language, Multisensory processing, Semantics, Auditory | 0-30: Visual, Language, Audition, Multisensory, Working memory, Cognitive control, Cued attention; 40-60: Emotion, Facial processing, Reward; 70-100: Motor, Pain, Autobiographical memory, Social cognition |

## Discussion

In this study we assessed the asynchronous connectivity of the entire brain during feature-length movie-watching, containing information that unfolds over a wide range of timescales, from momentary fluctuations in light and sound, to scenes of dialogue unfolding over minutes. We found that primary sensory regions receive the most numerous delayed inputs from the rest of the cortex (Figure 2E), over the longest (Figure 2A, B, D) and most variable timescales (Figure 2C), pointing towards a role as ‘hubs’ for asynchronous connections in the brain. We next applied two complementary dimensionality reduction techniques to investigate the organisation of asynchronous connectivity patterns. Using an unsupervised discrete-clustering approach, we found that the topographies delay maps could be meaningfully arranged into unique clusters of outputting seed regions that were distinguishable by their associated cognitive functions (Figure 3), thus indicating that the function of a given region is related to its delayed communications to the rest of the brain. Finally, using diffusion mapping to assess the organisation of delayed connectivity strength across the brain, we uncovered five whole-brain gradients of variation in intraregional reception of whole-brain connectivity. These gradients, each appear to be anchored by unique combinations of sensory and higher-order or subcortical regions (Figure 4). This suggests that the activities of both sensory and transmodal regions alike are influenced by multiple timescales of information, thus inviting a reevaluation of current hierarchical models of temporal processing in the brain.

### Whole brain delayed connectivity

The distribution of connectivity delays across the brain supports the placement of sensory processing regions at both the ‘start’ and ‘end’ of their respective sensory processing hierarchies. For instance, primary auditory cortices exhibited a median input delay of >2.1 seconds, out to >16 seconds at the 99th percentile of observed delays, indicating that these regions of the brain receive prior cortical information over word, sentence and discourse-level timescales. In contrast to traditional hierarchical brain models, which position them at the earliest^47^, and least temporally-integrated levels^8,15^, we consider our observation of sensory cortices as hubs for asynchronous inputs to be reflective of their putative demand for feedback of contextual information^5,22,48^. This may be indicative of processes by which longer-timescale (e.g., sentence-level) linguistic information ^1^ may shape ongoing activity of phonemic and word-level encoding processes in the auditory cortex, and other early sensory regions.

We interpret these results as being commensurate with a predictive processing framework. Therein, both descending predictions about upcoming stimuli and ascending signals encoding prediction error signals should both be received at their respective termini after some delay^49^. For example, a signal encoding a prediction of an upcoming word should not yield a prediction error signal from its terminus until at least the first phoneme of the predicted word has been heard (or not). Sensory regions receive a high degree of delayed inputs, largely because they are the recipients of substantial feedback from across the brain. This feedback, which carries predictions based on our previous experiences and current beliefs, helps to shape and refine sensory processing by continuously comparing these predictions with incoming sensory data.

Furthermore, the rate of transmission of descending predictive signals to their termini is limited by the temporal frequency of the information that they are representing^3,16^. Therefore, we might predict that delayed connectivity between regions should be reflective of particular temporal properties of ongoing stimulation. Indeed, we observed changes in connectivity delay between the first and final 20-minute segments of movies, which may reflect much longer-term tuning of asynchronous communication by narrative/contextual information (See Supplementary Figure S1). Therefore, the observed variation in the range of delays in communication between regions may facilitate appropriate scheduling of descending predictions that pertain to different timescales of information.

In addition to input delay maps, we computed approximations of the distribution of delays over which each region outputs connections to the rest of the cortex. These demonstrated a similar pattern; with the highest output-delays being observed in early sensory regions (Supplementary Figure 1), which we view to be complementary to the feedforward hierarchy of temporal receptivity put forward by Hasson^8^ et al. Specifically, while in this model, early sensory regions are most sensitive to rapid fluctuations in information encoded in sensory inputs, the brain-wide, connectivity-driven consequences of such sensitivity may occur over the longest timescales so as to facilitate the contextual accumulation of information, working memory, and other mnemonic processes^50^.

In reality, asynchronous communication between regions is likely to vary continuously with evolving contextual demands, as partially evidenced by intraregional differences in connectivity delay observed between earlier and later time points across movies (Supplementary Figure S2). Consequently, the prevalence of delayed connectivity observed across the brain warrants some speculation towards candidate mechanisms via which connection delays are incurred. Recurrent or re-entrant signalling, in principle, is defined by reciprocal connections between brain regions, allowing for sustained maintenance, amplification or attenuation of a particular signal over time^52,53^. Such signalling is highly prevalent across both local, and interregional circuits across the neocortex, and is considered to underpin the synchronisation and integration of cross-modal patterns of activity across the cortex^51^. Recently, the roles of locally reentrant circuits have been observed to underpin the ongoing maintenance of the ‘temporal pooling’ characteristics of neuronal populations^3,11,17,32^, which relates to the specific duration of time over which neural populations are sensitive to prior stimulus features^15,54,55^. Reentrant circuits may also underpin higher cortical functions, such as the guiding of attention, in the maintenance working memory^5150,56^, and the emergence of consciousness^27,57,58^.

As such, in the context of delayed connectivity between regions, the cross correlation function of these regions may be representative of the envelope of reentrant signalling process, over which a given signal is progressively amplified via fast cortico-thalamo-cortico signalling^59^ prior to decaying. The amplified signal pertaining to the representation of some stimulus feature from the recent past may then be leveraged for forthcoming predictions of stimulus input. Therefore, global reentrant activity may play a role in the shaping of the ‘temporal pooling’ of receptivity to prior cortical information^54^, such as the delayed input topography evidenced in the present study. This informs a novel hypothesis that the accurate ‘prediction’ of sensory information within primary sensory regions over the finest temporal resolutions demands the integration of information across longer, more diverse timescales. Consequently, the intra-regional tuning to time-windows of stimulus features of specific lengths^8^ may be supported by the delayed connections they receive from regions tuned to differing temporal windows^18,32^.

### Organisational principles of asynchronous connectivity

We also found that the topography of every unique delay map could be meaningfully organised into unique clusters of outputting seed regions. Specifically, our analysis revealed clusters of input-delay maps originating from both modality-specific, subcortical, and canonical resting-state network-specific seed regions; including frontoparietal and default mode networks. Separable networks of sensory and higher-order networks of regions were found to transmit connections over long and diverse timescales (Figure 3A), complementing the contemporary support for a feed-forward hierarchy of timescales across the cortical axis^11,20^, and feedback from transmodal regions^3,22,60^. This may represent a principal of network organisation in the brain by which information from functionally distinct networks of regions can be appropriately integrated elsewhere. For instance, signals output from subcortical regions (Figure 3.G) appear to be preferentially received by the rest of the brain over relatively short delays, allowing for rapid integration of bottom-up inputs. This observation is commensurate with contemporary understandings of the role of the thalamus as a coordinator of cortical information^61,62^. Being densely and recursively connected with every region of the cortex^63^, a core role of the thalamus lies in the synchronisation of salient information between cortical networks^64^. The low input and output delays of the thalamus observed in this study indicates the existence of ‘fast transfer’ networks, where cortical information can be integrated, normalised and amplified before being relayed to other networks. Taken together, these groupings signify functional differentiation between, and various modes of non-simultaneous communication within, distinct networks of outputting regions. As the networks that emerged from this data-driven approach respectively appeared to be associated with separable perceptual and higher order modalities, it may be that the functional specialisation of regions is supported by the topography of their asynchronous connectivity to the rest of the brain^65^.

However, this discrete network perspective overlooks the overlapping roles of individual regions across distributed networks supporting various perceptual and cognitive faculties^5^. Instead, each network is likely to dynamically engage with others according to task demands^15,16,66,67^. Hence, in a complementary analysis, we hypothesised that close associations between sensory and higher-order regions would be observed along orthogonal gradients of delayed connectivity, as opposed to connection delays, reflecting their respective roles as sites integration of temporally extended information. We uncovered five gradients of variation in received delayed connectivity across the brain, anchored by unique combinations of sensory and higher-order or subcortical regions.

This finding contrasts with conventional hierarchical views that depict a feedforward accumulation of information along the cortical hierarchy^11,15,20^. Rather than suggesting that higher-order processing networks remain uniformly at a maximum distance from sensory regions^21^, instead, this points towards multiple modes of specific, temporally extended pairings between certain networks associated with sensory modalities and higher-order functions that are otherwise obscured by instantaneous measures of functional connectivity. Specifically, modality-specific combinations of higher-order and early sensory regions receive similar sets of asynchronous connections from the rest of the brain, suggesting that predictive hierarchies aren’t necessarily organised along a gradient of perception to cognition, but may rather be dynamically reconfigurable according to task demands^5,68,69^. This *heterarchical*^41^ pattern of associations is also commensurate with the brain’s ability to integrate and process, temporally extended multimodal information within separable ‘modes of operation’ within an ontology of cognition that doesn’t arbitrarily draw distinctions between perception and higher-order functions. For instance, gradient five (Figure 4F) appears to separate regions associated with the topics relating to empathy, including ‘pain’, ‘motor’ and ‘social cognition’, from a network of regions associated with visual problem-solving, including ‘visual semantics’, ‘numerical cognition’ and ‘eye movements’.

### Limitations

The impact of heterogeneity of neurovascular coupling speeds across the brain^70^ on the measurement of functional connectivity is double-edged. Given the slowly-unfolding nature of the haemodynamic response, the peak of which lasts for up to two seconds^71^, standard measures of functional connectivity may mischaracterize veridically delayed correlations as instantaneous. That said, differences in the time-to-peak hemodynamic response function (HRF) between regions may account for some variance in connectivity delays between regions, although it is highly unlikely that this would extend to the >70 second delays we observed between some regions (Table 1). While the potential influence of the HRF on how connectivity delays are distributed across the brain was not considered in our analyses, previous studies investigating asynchronous connectivity have attributed observed latencies to neural activity, rather than being solely hemodynamic in nature^72,73^, and the spatial distribution of time-to-peak HRF overlaps only minimally with the topography of delayed connectivities observed in the present study^74,75^.

A further limitation of the current work also plagues the majority of investigations of functional connectivity in the brain, in that it fails to account for non-linear interactions between regions ^76^, or multivariate synergies and redundancies in information shared between regions ^77^. For instance, Varley et al recently demonstrated that instantaneous correlations were extremely highly correlated (R > 0.999, p :S 10^-20^) with redundant information in an analysis of multivariate relationships between the activity timeseries of brain regions ^78^. Despite offering clear advantages over instantaneous measures of functional connectivity with our detection of directed linear, positive relationships between brain regions, our measure of delayed connectivity may still largely capture redundant information, especially in cases where ‘optimal’ communication paths between regions in terms of shortest delay-paths may first flow through lower-latency regions ^73,79^. Future investigations of the asynchronous communication in the brain could therefore benefit by considering non-linear, anti-correlated and multivariate interactions between regions.

## Conclusion

We have presented a novel technique; ‘delayed connectivity mapping’, with the goal of providing evidence towards a model of cognition across timescales that unifies distributed, predictive processing accounts of brain function^22^ with extant evidence for functional specialisation in the processing of multiple timescales of information^11^. Specifically, the ability of early sensory cortices to preferentially track rapid fluctuations in sensory information is likely facilitated by their receipt of extensive, asynchronous feedback from the rest of the brain as evidenced in this study. Latent structure in the spatial distribution of delayed connections between regions further supported an association between asynchronous connectivities and intra-regional functional specialisation. Here, using an unsupervised clustering approach, we revealed distinct clusters of functionally-related brain regions characterised by their delayed communication patterns, indicating that functional specialisation is associated with the timing and organisation of asynchronous connections within the brain. In a complementary approach, separable gradients of variability in the received connections of regions reveal distinct sets of similarities between sensory, and transmodal networks, revealing overlapping modes of communication over extended time frames between these sets of regions. These results challenge contemporary conceptions of the temporal hierarchy within the brain by emphasising the role of sensory regions as sites of temporal integration across timescales. Taken together, our results also support an integrative model of the human brain connectome, wherein regional functional specialisation is supported by asynchronous, distributed processing^65,80,81^.

## Methods

### Participants

The present study analysed the ‘Naturalistic Neuroimaging Database’ (NNDb)^33^. The NNDb consists of 86 participants undergoing fMRI scans while watching one of 10 previously unseen full-length movies selected across a range of genres. All participants were right handed native English speakers with no MRI contraindications, no visual or hearing impairments and no current use of prescribed medication. Participants’ ages ranged from 18-58, and the sample consisted of 42 females / 44 males. All participants provided written informed consent to take part in the study and share their anonymised data. Full study procedures and participant demographics are available on https://www.naturalistic-neuroimaging-database.org/.

### MRI Acquisition

Functional and anatomical images were acquired using 1.5T Siemens MAGNETOM Avanto with a 32 channel head coil (Siemens Healthcare, Erlangen, Germany). A multiband EPI sequence was employed^82,83^, using the following parameters: TR = 1 s TE = 54.8 ms, a flip angle of 75°, 40 interleaved slices, an isotropic resolution of 3.2 mm with a 4x multiband factor and no in-plane acceleration; to reduce cross-slice aliasing^84^, the ‘leak block’ option was enabled^85^. This EPI sequence had a software limitation of one hour of consecutive scanning, meaning each movie had at least one break. One or two cerebellar slices were missing from participants with large heads. Anatomical scans were acquired after functional scans, involving a 10 min high-resolution T1-weighted MPRAGE (TR = 2.73 s, TE = 3.57 ms, 176 sagittal slices, resolution = 1.0 mm).

During functional scans, participants were presented with feature length movie stimuli presented on a mirrored, back-projected 22.5 x 42cm screen, at a viewing angle of 19.0°. The screen was positioned behind the head coil and was viewed through a mirror placed above participants’ eyes. Audio was presented in stereo via MRI compatible, scanner noise-attenuating headphones

### Preprocessing

Anatomical images were de-skulled and non-linearly aligned to the *MNT152_2009_template_SSW.nii.gz* template brain^86^ using the default settings within AFNI’s @SSWarper tool, prior to inflation and registration using default settings in Freesurfers ‘recon-all’ tool (V6.0)^87^. Automated anatomical parcellations generated during the recon-all step were then used to calculate white-matter and ventricle Regions of Interest (ROIs), and subsequently eroded to exclude any grey matter voxels, for later use as noise regressors.

Next, the standardised pipeline, afni_proc.py was used to preprocess all functional images. All parameters for the pre-processing pipeline are available at https://openneuro.org/datasets/ds002837/versions/2.0.0. All images were de-spiked and corrected for slice timing differences. Participants’ EPI volumes were registered to the statistically determined lowest-motion volume (volreg_align_to MIN_OUTLIER), and aligned to their corresponding anatomical images (volreg_align_e2a). These volume-registered, anatomically aligned time series were nonlinearly aligned to the MNI template brain (volreg_align_tlrc). Time Series were cleaned and normalised using parameters accessible on the NNDb openneuro repository (https://openneuro.org/datasets/ds002837/versions/2.0.0)

All detrended time series were next concatenated, on which spatial ICA was then performed over 250 dimensions using FSL’s melodic tool^88^ for removal of artefacts and to increase signal to noise ratio^89^. Nuisance regressors were manually selected and regressed out of the detrended, concatenated time series (see)^33^.

Two versions of the pre-processed, detrended and concatenated time series with ICA-based artefacts removed were generated: one with spatial blurring and one without. The blurred timeseries were used for all further analysis, with the exception of functional-anatomical parcellation steps outlined below, in order to account for overlapping boundaries between functionally discrete regions.

### Delay Mapping

#### Region of Interest parcellation and signal extraction

In order to account for ‘soft’ borders and overlaps between anatomical ROIs and preserve functional gradients in signal within each region ^20^, we employed the Difumo functional parcellation and signal extraction technique^90^. This yielded 1024 ROI timeseries covering the entire brain for each participant, at each sampling time point from unblurred, uncensored time series. All Difumo ROIs labelled as predominantly containing cerebrospinal fluid and white matter were manually discarded, leaving a total of 807 ROIs for further analysis.

#### Delay estimation

Time-delay between each pair of reference time series and voxel time series from the rest of the brain was then estimated using a computationally efficient algorithm as implemented by 3dDelay in AFNI, forcing >0 delay values into each voxel. In this context, time-delay was determined via estimation of peak of the cross-correlation function between reference and voxel-wise time series.

In principle, this is achieved by calculating the Pearson’s correlation coefficient at systematically introduced lags between reference time series and voxel time series to produce the cross-correlation function. It naturally follows that ‘delay’ is simply the maximum point of the cross-correlation function. As fMRI time series are discrete series, the estimation of delay at sub-TR resolutions requires interpolation about maximal points in the calculated cross-correlation function. As detailed by Saad, DeYoe and Ropella (2003), the algorithm employed by 3dDelay efficiently estimates delay to a sub-TR resolution via linear interpolation about 0 of the Hilbert transform of the envelope of the cross correlation function calculated between ROI and voxel time series.

Using this method, ROI-wise estimates of voxel-wise delays were generated and will henceforth be referred to as ‘ROI delay maps’. It is also important to note at this point that the above method for estimating delay will always generate non-integer estimates of values greater than zero, within every voxel. Voxels present within the ROI used to generate each ROI delay map were set to zero in order to remove artefacts resulting from intra-regional autocorrelations. As 3dDelay outputs the delay at which cross-correlation is maximal between regions regardless of the strength of correlation at maximal lags, it was necessary to threshold ROI delay maps by a minimum cross-correlation coefficient of 0.1 in order to remove spurious delayed connectivities.

#### Delayed Global Input estimation

Across each of the ROI-wise delay maps generated in the previous step, any given voxel of the brain will exhibit a distribution of delay values, representing the range of timescales over which the voxel receives inputs from the rest of the cortex. In order to characterise useful descriptive features of this distribution, we calculated the mean, median, 99th percentile and standard deviation across all ROI-wise delay maps for each non-zero voxel in the brain. Additionally, we record the total count of above-threshold ROI-wise delay maps that contributed to these statistics in each voxel (henceforth, input count; representing the ‘temporal centrality’ of a given region, or its propensity to receive cortical inputs across observed delays), as well as the maximum value observed across each contributing map.

#### Delayed Global Output estimation

Within each participant, the mean across all non-zero voxels of each thresholded delay map is calculated to provide an estimation of the delayed global output of each ROI. A binary mask of each ROI is then multiplied by their corresponding delayed global output. To achieve smooth boundaries between overlapping DiFumo ROIs, the mean of each of these ROI output-maps across non-zero voxels is calculated to yield maps of the average delayed output of each region in one volume per participant. This process was repeated to generate output maps for the median, standard deviation and 99th percentile of global delays

#### Group-level analysis

Group analysis of each delayed input and output map was performed via linear mixed effects modelling as implemented by 3dLME in AFNI^91^. Movie, age and gender were included as fixed effects in each identical model, while intra-participant variation in voxel intensity was modelled as a random effect.

It is initially unclear as to whether whole-brain delayed connectivity estimation reflects an intrinsic property of asynchronous functional connectivity under naturalistic conditions, or whether delays between regions reflect properties of the stimulus. Should they reflect stimulus properties, some changes in delays should be observable as contextual information in the movie accumulates. In order to do so, 20-minute movie segments (taken from the beginning, middle and end of movies) were also included as fixed effects in the model, and post-hoc general linear tests were computed for direct contrasts between the factor level effects of these segments.

To correct for multiple comparisons in all group-level analyses across both datasets, a minimum cluster size was determined from voxel-wise p-value thresholds such that any clusters over a given size would reach a corrected threshold of alpha = 0.01. This was implemented via the 3dClustsim tool in AFNI, which calculates the probability of false-positive of clusters in a null model generated by Monte Carlo simulations (set to 10000 permutations) of Gaussian noise and the spatial autocorrelation of residuals generated in each group-level analysis. Cluster sizes (Detection Threshold < 0.01) were estimated across voxel-wise p-values of 0.05, 0.02, 0.01, 0.005, 0.002, 0.001 prior to thresholding of each volume of interest at corresponding z-values for each p-value and its associated cluster size before finally merging resulting volumes. Surviving clusters were projected onto a standard MNI surface model of the brain and visualised in BrainNet Viewer ^92^.

#### Recovery of modality-specific networks from output-delay topography

The above technique for calculating estimates for the voxel-wise aggregate statistics for delayed input maps are likely to obscure meaningfully separable spatial patterns of delayed input connectivity generated by individual ROIs.

To do so, group (median) averages of each un-threshold ROI-wise delay map were computed across participants, movies, and movie segments, yielding 807 ROI-wise average delay maps. From each volume, delay values were extracted from each voxel using *3dMaskDump* to form a vector of delay values. A negative distance matrix was generated, representing geodesic (Manhattan) distances between each vector was computed, where each value represents a (dis)similarity score for the spatial distributions of delay values across the cortex, between ROI-wise maps. Column-wise clustering of group-average delay maps was then performed in R, using the *APCluster* library^93^. Hierarchical clusters were then determined with the default input preference of q=0.5, corresponding to the median of input similarities.

The ROI-wise, group average delayed connectivity maps comprising each cluster revealed via this method were then averaged, generating cluster-specific delay-topographies. The outputting Difumo ROIs comprising each cluster were also combined into a single volume to generate estimates of the functional network giving rise to each cluster’s delay topography. Superimposition of the outline of outputting regions over cluster delay maps was implemented using the GIMP image editing software.

#### Spatial gradients of delayed connectivity

ROI-wise cross-correlation coefficient maps (CCMs) were first extracted from volumes output by *3dDelay* in the aforementioned ‘Delay Estimation’ step. Each voxel in these maps is assigned a value corresponding to the peak of the cross-correlation coefficient between reference ROI and target voxel time series, yielding a measure of connectivity-at-optimal-delay. We then calculated a group-level average of each map, yielding 807 ROI-wise CCMs. We then extracted the average value within binary masks of each of the 807 DiFumo ROIs from each of the 807 CCMs using *3dMaskAve*, yielding a ROI x ROI asymmetric directed connectivity-at-optimal-delay matrix which was then normalised.

Diffusion mapping was implemented via default parameters (10 components computed with the diffusion mapping approach over a cosine similarity matrix thresholded to 5% sparsity) provided by the *’GradientMaps’* function in the BrainSpace toolbox ^38^ to remove potential negative similarities and ensure only the least noisy connections are considered in the embedding solution. As the delay-adjusted connectivity matrix was asymmetric, this procedure was performed again on a 90° rotation of the original matrix, yielding one set of gradients for ‘input’ connectivities, and another for ‘outputs’. Output gradients are not presented as they failed to reach the eigenvalue threshold of >1.

To represent how these ‘gradients of delayed connectivity’ vary across the cortex, embeddings for principal component embedding values for each column vector in the affinity matrix (i.e., x and y coordinates, respectively) were projected back into the voxels comprising the DiFumo parcel from which the delay map associated with the vector was originally generated. The fuzzy boundaries of DiFumo parcellations allow for continuous representation of ‘delayed connectivity gradients’ across the cortex.

#### Neurosynth Decoding

Interpretation of both whole-brain topographic activity maps, and ‘blob-wise’ contrasts are open to bias in terms of the functional assignments of given regions. In order to mitigate such bias, across appropriate analyses, we employed two separate decoding methods, both working via comparison with the Neurosynth Database (Version 0.7, https://github.com/neurosynth/neurosynth-data; Yarkoni, Poldrack, Nichols, Van Essen, & Wager, 2011). The database contains over 500,000 activation peaks or centers of mass collected across 14,371 studies with over 3200 term-based features, from which neurosynth can rapidly synthesise term-based meta-analytic probabilistic maps. Decoding of analysis volumes is then achieved by systematically examining the spatial correlation between input volumes and meta-analytic maps.

We first sought to assign functions to ROIs recovered from our aforementioned clustering method (see section titled ‘Recovery of Delayed Connectivity Networks’). To do so, we employed the ‘DiscreteROIAssociation’ decoding method as implemented in the NiMare library for Python (Salo et al., 2021) to decode ‘outputting’ ROIs recovered from our clustering analysis. This method compares discrete ROIs to the entire Neurosynth (Version 0.7) database, for which we present the top 50 terms for each cluster in wordcount format in Figure 2 and supplementary Figures S4, S5.

Additionally, we implemented a previously employed topic-wise decoding method to find the association between each of 24 topics and respective cluster-wise, and diffusion gradient-percentile maps. This method utilised code that was modified from an open-source repository available at https://www.github.com/gpreti/GSP_StructuralDecouplingIndex, run in a custom python2.7 environment. The topics were derived from previous research ^19,94,95^ and include ‘lower order’ sensory functions such as audition, visual perception, eye movement, motion, to higher order features including working memory, numerical cognition, attention, affect and working memory. Maps of the ‘outputting regions’ that generate each cluster revealed by APC were used for the topic-association analysis presented in Figure 2, while binarised masks of five-percentile bins ranging from the lowest 5th percentile to the highest 95th percentile of gradient maps were used for the analysis presented in Figure 3. In all cases, topic-terms were ordered according to the weighted mean of the resulting decoding z-statistics.

## References

1. Stephens, G. J., Honey, C. J. & Hasson, U. A place for time: the spatiotemporal structure of neural dynamics during natural audition. J. Neurophysiol. 110, 2019–2026 (2013).

2. Baldassano, C. et al. Discovering Event Structure in Continuous Narrative Perception and Memory. Neuron 95, 709–721.e5 (2017).

3. Kiebel, S. J., Daunizeau, J. & Friston, K. J. A hierarchy of time-scales and the brain. PLoS Comput. Biol. 4, e1000209 (2008).

4. Skipper, J. I. Echoes of the spoken past: how auditory cortex hears context during speech perception. Philos. Trans. R. Soc. Lond. B Biol. Sci. 369, 20130297 (2014).

5. Skipper, J. I. The NOLB model: a model of the natural organization of language and the brain. in Cognitive Neuroscience of Natural Language Use (ed. Willems, R. M.) 101–134 (Cambridge University Press, 2015).

6. Skipper, J. I., Devlin, J. T. & Lametti, D. R. The hearing ear is always found close to the speaking tongue: Review of the role of the motor system in speech perception. Brain Lang. 164, 77–105 (2017).

7. Ulanovsky, N., Las, L., Farkas, D. & Nelken, I. Multiple time scales of adaptation in auditory cortex neurons. J. Neurosci. 24, 10440–10453 (2004).

8. Hasson, U., Yang, E., Vallines, I., Heeger, D. J. & Rubin, N. A hierarchy of temporal receptive windows in human cortex. J. Neurosci. 28, 2539–2550 (2008).

9. Lerner, Y., Honey, C. J., Silbert, L. J. & Hasson, U. Topographic mapping of a hierarchy of temporal receptive windows using a narrated story. J. Neurosci. 31, 2906–2915 (2011).

10. Farbood, M. M., Heeger, D. J., Marcus, G., Hasson, U. & Lerner, Y. The neural processing of hierarchical structure in music and speech at different timescales. Front. Neurosci. 9, 157 (2015).

11. Hasson, U., Chen, J. & Honey, C. J. Hierarchical process memory: memory as an integral component of information processing. Trends Cogn. Sci. 19, 304–313 (2015).

12. Murray, J. D. et al. A hierarchy of intrinsic timescales across primate cortex. Nat. Neurosci. 17, 1661–1663 (2014).

13. Cavanagh, S. E., Wallis, J. D., Kennerley, S. W. & Hunt, L. T. Autocorrelation structure at rest predicts value correlates of single neurons during reward-guided choice. Elife 5, (2016).

14. Watanabe, T., Rees, G. & Masuda, N. Atypical intrinsic neural timescale in autism. Elife 8, (2019).

15. Golesorkhi, M., Gomez-Pilar, J., Tumati, S., Fraser, M. & Northoff, G. Temporal hierarchy of intrinsic neural timescales converges with spatial core-periphery organization. Commun Biol 4, 277 (2021).

16. Gollo, L. L., Zalesky, A., Hutchison, R. M., van den Heuvel, M. & Breakspear, M. Dwelling quietly in the rich club: brain network determinants of slow cortical fluctuations. Philos. Trans. R. Soc. Lond. B Biol. Sci. 370, (2015).

17. Wolff, A. et al. Intrinsic neural timescales: temporal integration and segregation. Trends Cogn. Sci. 26, 159–173 (2022).

18. Chaudhuri, R., Knoblauch, K., Gariel, M.-A., Kennedy, H. & Wang, X.-J. A Large-Scale Circuit Mechanism for Hierarchical Dynamical Processing in the Primate Cortex. Neuron 88, 419–431 (2015).

19. Margulies, D. S. et al. Situating the default-mode network along a principal gradient of macroscale cortical organization. Proc. Natl. Acad. Sci. U. S. A. 113, 12574–12579 (2016).

20. Huntenburg, J. M., Bazin, P.-L. & Margulies, D. S. Large-Scale Gradients in Human Cortical Organization. Trends Cogn. Sci. 22, 21–31 (2018).

21. Samara, A., Eilbott, J., Margulies, D. S., Xu, T. & Vanderwal, T. Cortical gradients during naturalistic processing are hierarchical and modality-specific. Neuroimage 271, 120023 (2023).

22. Clark, A. Whatever next? Predictive brains, situated agents, and the future of cognitive science. Behav. Brain Sci. 36, 181–204 (2013).

23. Kayser, C., Kording, K. P. & Konig, P. Processing of complex stimuli and natural scenes in the visual cortex. Curr. Opin. Neurobiol. 14, 468–473 (2004).

24. Petro, L. S., Vizioli, L. & Muckli, L. Contributions of cortical feedback to sensory processing in primary visual cortex. Front. Psychol. 5, 1223 (2014).

25. Budinger, E. & Scheich, H. Anatomical connections suitable for the direct processing of neuronal information of different modalities via the rodent primary auditory cortex. Hear. Res. 258, 16–27 (2009).

26. Winer, J. A. & Schreiner, C. E. The auditory cortex. (2010).

27. Edelman, G. M. The remembered present: a biological theory of consciousness. J. Cogn. Neurosci. 2, 385–387 (1990).

28. Grill-Spector, K. & Malach, R. The human visual cortex. Annu. Rev. Neurosci. 27, 649–677 (2004).

29. Hsieh, P.-J., Vul, E. & Kanwisher, N. Recognition alters the spatial pattern of FMRI activation in early retinotopic cortex. J. Neurophysiol. 103, 1501–1507 (2010).

30. Gonzalez-Garcia, C. & He, B. J. A Gradient of Sharpening Effects by Perceptual Prior across the Human Cortical Hierarchy. J. Neurosci. 41, 167–178 (2021).

31. Hedger, N. & Knapen, T. Naturalistic Audiovisual Stimulation Reveals the Topographic Organization of Human Auditory Cortex. bioRxiv 2021.07.05.447566 (2021) doi:10.1101/2021.07.05.447566.

32. Golesorkhi, M. et al. The brain and its time: intrinsic neural timescales are key for input processing. Commun Biol 4, 970 (2021).

33. Aliko, S., Huang, J., Gheorghiu, F., Meliss, S. & Skipper, J. I. A ‘Naturalistic Neuroimaging Database’ for understanding the brain using ecological stimuli. bioRxiv 2020.05.22.110817 (2020) doi:10.1101/2020.05.22.110817.

34. Sonkusare, S., Breakspear, M. & Guo, C. Naturalistic Stimuli in Neuroscience: Critically Acclaimed. Trends Cogn. Sci. 23, 699–714 (2019).

35. Finn, E. S. Is it time to put rest to rest? Trends Cogn. Sci. 25, 1021–1032 (2021).

36. Barron, H. C., Garvert, M. M. & Behrens, T. E. J. Repetition suppression: a means to index neural representations using BOLD? Philos. Trans. R. Soc. Lond. B Biol. Sci. 371, (2016).

37. Saad, Z. S., DeYoe, E. A. & Ropella, K. M. Estimation of FMRI response delays. Neuroimage 18, 494–504 (2003).

38. Vos de Wael, R., et al. BrainSpace: a toolbox for the analysis of macroscale gradients in neuroimaging and connectomics datasets. Commun Biol 3, 103 (2020).

39. Pessoa, L. A Network Model of the Emotional Brain. Trends Cogn. Sci. 21, 357–371 (2017).

40. Spagna, A., et al. Heterarchy in Visual Mental Imagery: a review of methods and neurobehavioral findings. (2023).

41. McCulloch, W. S. A heterarchy of values determined by the topology of nervous nets. Bull. Math. Biophys. 7, 89–93 (1945).

42. Frey, B. J. & Dueck, D. Clustering by passing messages between data points. Science 315, 972–976 (2007).

43. Salo, T. et al. NiMARE: Neuroimaging meta-analysis research environment. NeuroLibre 1, 7 (2022).

44. Ito, T., Hearne, L. J. & Cole, M. W. A cortical hierarchy of localized and distributed processes revealed via dissociation of task activations, connectivity changes, and intrinsic timescales. bioRxiv 262626 (2020) doi:10.1101/262626.

45. Mckeown, B. et al. The relationship between individual variation in macroscale functional gradients and distinct aspects of ongoing thought. Neuroimage 220, 117072 (2020).

46. Marek, S. & Dosenbach, N. U. F. The frontoparietal network: function, electrophysiology, and importance of individual precision mapping. Dialogues Clin. Neurosci. 20, 133–140 (2018).

47. Mesulam, M. M. From sensation to cognition. Brain 121 **(Pt** **6**), 1013–1052 (1998).

48. Hochstein, S. & Ahissar, M. View from the top: hierarchies and reverse hierarchies in the visual system. Neuron 36, 791–804 (2002).

49. Friston, K. Learning and inference in the brain. Neural Netw. 16, 1325–1352 (2003).

50. Voitov, I. & Mrsic-Flogel, T. D. Cortical feedback loops bind distributed representations of working memory. Nature 608, 381–389 (2022).

51. Edelman, G. M. & Gally, J. A. Reentry: a key mechanism for integration of brain function. Front. Tntegr. Neurosci. 7, 63 (2013).

52. Douglas, R. J. & Martin, K. A. C. Neuronal circuits of the neocortex. Annu. Rev. Neurosci. 27, 419–451 (2004).

53. Rutishauser, U. & Douglas, R. J. State-dependent computation using coupled recurrent networks. Neural Comput. 21, 478–509 (2009).

54. Himberger, K. D., Chien, H.-Y. & Honey, C. J. Principles of Temporal Processing Across the Cortical Hierarchy. Neuroscience 389, 161–174 (2018).

55. Beiran, M. & Ostojic, S. Contrasting the effects of adaptation and synaptic filtering on the timescales of dynamics in recurrent networks. PLoS Comput. Biol. 15, e1006893 (2019).

56. Wang, X. J. Synaptic basis of cortical persistent activity: the importance of NMDA receptors to working memory. J. Neurosci. 19, 9587–9603 (1999).

57. Tsuchiya, N. & van Boxtel, J. J. A. Is recurrent processing necessary and/or sufficient for consciousness? Cogn. Neurosci. 1, 230–231 (2010).

58. Mashour, G. A., Roelfsema, P., Changeux, J.-P. & Dehaene, S. Conscious Processing and the Global Neuronal Workspace Hypothesis. Neuron 105, 776–798 (2020).

59. Wolff, M., Morceau, S., Folkard, R., Martin-Cortecero, J. & Groh, A. A thalamic bridge from sensory perception to cognition. Neurosci. Biobehav. Rev. 120, 222–235 (2021).

60. Caucheteux, C., Gramfort, A. & King, J.-R. Evidence of a predictive coding hierarchy in the human brain listening to speech. Nat Hum Behav (2023) doi:10.1038/s41562-022-01516-2.

61. Steriade, M. & Pare, D. Gating in Cerebral Networks. (Cambridge University Press, 2007).

62. Moustafa, A. A., McMullan, R. D., Rostron, B., Hewedi, D. H. & Haladjian, H. H. The thalamus as a relay station and gatekeeper: relevance to brain disorders. Rev. Neurosci. 28, 203–218 (2017).

63. Jones, E. G. The Thalamus. (Springer Science & Business Media, 2012).

64. Wang, Q., Webber, R. M. & Stanley, G. B. Thalamic synchrony and the adaptive gating of information flow to cortex. Nat. Neurosci. 13, 1534–1541 (2010).

65. Pessoa, L. Understanding brain networks and brain organization. Phys. Life Rev. 11, 400–435 (2014).

66. Skipper, J. I. A voice without a mouth no more: The neurobiology of language, consciousness, and mental health. (2021) doi:10.31234/osf.io/jfuws.

67. Klar, P., Çatal, Y., Langner, R., Huang, Z. & Northoff, G. Scale-free dynamics of core-periphery topography. Hum. Brain Mapp. 44, 1997–2017 (2023).

68. Simony, E. et al. Dynamic reconfiguration of the default mode network during narrative comprehension. Nat. Commun. 7, 12141 (2016).

69. Aliko, S., Wang, B., Small, S. L. & Skipper, J. I. The entire brain, more or less is at work: ‘Language regions’ are artefacts of averaging. bioRxiv 2023.09.01.555886 (2023) doi:10.1101/2023.09.01.555886.

70. Handwerker, D. A., Ollinger, J. M. & D’Esposito, M. Variation of BOLD hemodynamic responses across subjects and brain regions and their effects on statistical analyses. Neuroimage 21, 1639–1651 (2004).

71. Mayer, A. R. et al. Investigating the properties of the hemodynamic response function after mild traumatic brain injury. J. Neurotrauma 31, 189–197 (2014).

72. Mitra, A., Snyder, A. Z., Hacker, C. D. & Raichle, M. E. Lag structure in resting-state fMRI. J. Neurophysiol. 111, 2374–2391 (2014).

73. Mitra, A., Snyder, A. Z., Blazey, T. & Raichle, M. E. Lag threads organize the brain’s intrinsic activity. Proc. Natl. Acad. Sci. U. S. A. 112, E2235–44 (2015).

74. Wu, G.-R. et al. A blind deconvolution approach to recover effective connectivity brain networks from resting state fMRI data. Med. Tmage Anal. 17, 365–374 (2013).

75. Taylor, A. J., Kim, J. H. & Ress, D. Characterization of the hemodynamic response function across the majority of human cerebral cortex. Neuroimage 173, 322–331 (2018).

76. Motlaghian, S. M. et al. Nonlinear functional network connectivity in resting functional magnetic resonance imaging data. Hum. Brain Mapp. 43, 4556–4566 (2022).

77. Varley, T. F. & Sporns, O. Network Analysis of Time Series: Novel Approaches to Network Neuroscience. Front. Neurosci. 15, (2022).

78. Varley, T. F., Pope, M., Maria Grazia, Joshua & Sporns, O. Partial entropy decomposition reveals higher-order information structures in human brain activity. Proc. Natl. Acad. Sci. U. S. A. 120, e2300888120 (2023).

79. Lizier, J. T., Bauer, F., Atay, F. M. & Jost, J. Analytic relationship of relative synchronizability to network structure and motifs. arXiv [cs.ST] (2023).

80. Anderson, M. L., Kinnison, J. & Pessoa, L. Describing functional diversity of brain regions and brain networks. Neuroimage 73, 50–58 (2013).

81. Sorrentino, P. et al. Whole-brain propagation delays in multiple sclerosis, a combined tractography - magnetoencephalography study. J. Neurosci. (2022) doi:10.1523/JNEUROSCI.0938-22.2022.

82. Feinberg, D. A. et al. Multiplexed echo planar imaging for sub-second whole brain FMRI and fast diffusion imaging. PLoS One 5, e15710 (2010).

83. Feinberg, D. A. & Setsompop, K. Ultra-fast MRI of the human brain with simultaneous multi-slice imaging. J. Magn. Reson. 229, 90–100 (2013).

84. Todd, N. et al. Evaluation of 2D multiband EPI imaging for high-resolution, whole-brain, task-based fMRI studies at 3T: Sensitivity and slice leakage artifacts. Neuroimage 124, 32–42 (2016).

85. Cauley, S. F., Polimeni, J. R., Bhat, H., Wald, L. L. & Setsompop, K. Interslice leakage artifact reduction technique for simultaneous multislice acquisitions. Magn. Reson. Med. 72, 93–102 (2014).

86. Holmes, C. J. et al. Enhancement of MR images using registration for signal averaging. J. Comput. Assist. Tomogr. 22, 324–333 (1998).

87. Fischl, B. FreeSurfer. Neuroimage 62, 774–781 (2012).

88. Smith, S. M. et al. Advances in functional and structural MR image analysis and implementation as FSL. Neuroimage 23 **Suppl 1**, S208–19 (2004).

89. Liu, X., Zhen, Z., Yang, A., Bai, H. & Liu, J. A manually denoised audio-visual movie watching fMRI dataset for the studyforrest project. Sci Data 6, 295 (2019).

90. Dadi, K. et al. Fine-grain atlases of functional modes for fMRI analysis. Neuroimage 221, 117126 (2020).

91. Chen, G., Saad, Z. S., Britton, J. C., Pine, D. S. & Cox, R. W. Linear mixed-effects modeling approach to FMRI group analysis. Neuroimage 73, 176–190 (2013).

92. Xia, M., Wang, J. & He, Y. BrainNet Viewer: a network visualization tool for human brain connectomics. PLoS One 8, e68910 (2013).

93. Bodenhofer, U., Kothmeier, A. & Hochreiter, S. APCluster: an R package for affinity propagation clustering. Bioinformatics 27, 2463–2464 (2011).

94. Luppi, A. I. et al. A synergistic core for human brain evolution and cognition. Nat. Neurosci. 25, 771–782 (2022).

95. Preti, M. G. & Van De Ville, D. Decoupling of brain function from structure reveals regional behavioral specialization in humans. Nat. Commun. 10, 4747 (2019).

