## Supplementary Information for "Where the present gets remembered: Sensory regions communicate with the brain over the longest timescales"

Output maps were generated by computing the global statistics (mean, median, standard deviation and 99th percentile) of each ROI-wise delay map, and projecting these values back into the voxels of each respective 'outputting' ROI. As compared to median input-delay maps (Figure 2A), delay values are longer, more variable (standard deviation), and more diffusely distributed in output delay maps, covering a large portion of the medial temporal and angular gyri, in addition to the superior and inferior parietal lobules. It is also noteworthy that the cerebellar crus II demonstrates comparatively long input and output delayed connections, while the cerebellar crus III shows long delayed outputs.

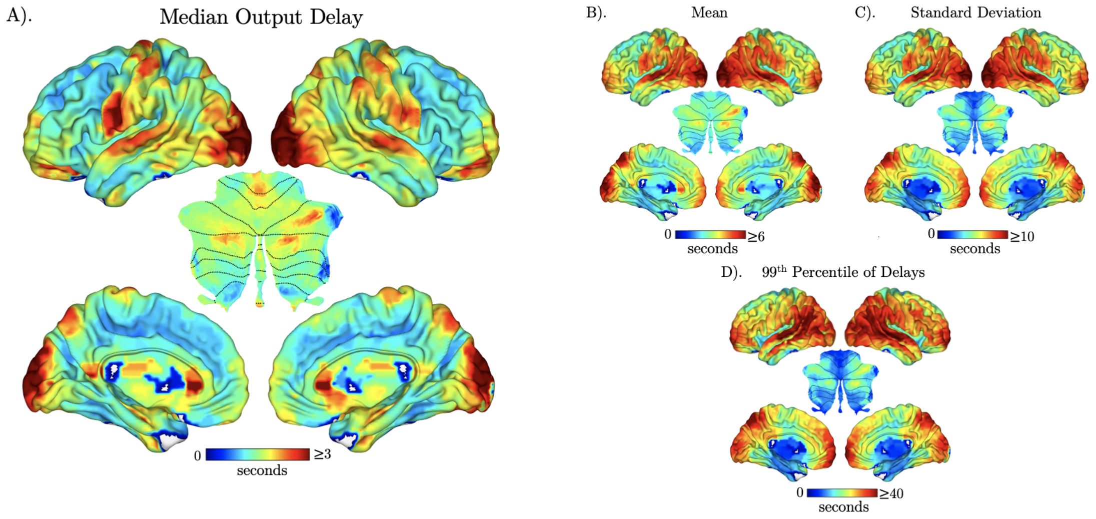


***Supplementary Figure S1)*** *Delayed Output Connectivity Maps generated using a group-level linear mixed effects model. The depicted statistics across three movie segments include: (A) median output delay in seconds, (B) mean, (C) standard deviation, (D) 99th percentile of output delays to the rest of the brain with cross-correlation coefficient $R \geq 0.1$. All outcomes are thresholded and corrected for multiple comparisons at alpha = 0.01.*

As connectivity delays between may covary with contextual information that accumulates over the course of a movie, we also assessed temporal stability of delayed connectivity topography throughout the films. We observed decreases in the median, mean and standard deviation of input delays received by occipital and parietal regions, as well as the posterior cingulate cortex and precuneus (Supplementary Figure S2.A.i,ii,iii,v, region-specific magnitude changes detailed in Supplementary Table T1). Conversely, widespread increases in the number of above-threshold inputs received were observed across regions including the supplementary motor area, precuneus, medial temporal lobe, superior, and middle temporal gyri. Similar stimulus-dependent changes between first and final chunks were observed in output-delay maps (Supplementary figure S1.B). Conversely, increases in the number of input-connections above threshold (R≥0.1) were observed in superior and medial temporal temporal gyri, middle frontal gyri, thalamus, caudate nucleus, and lingual and fusiform gyri (Supplementary Figure S1.A.iv).

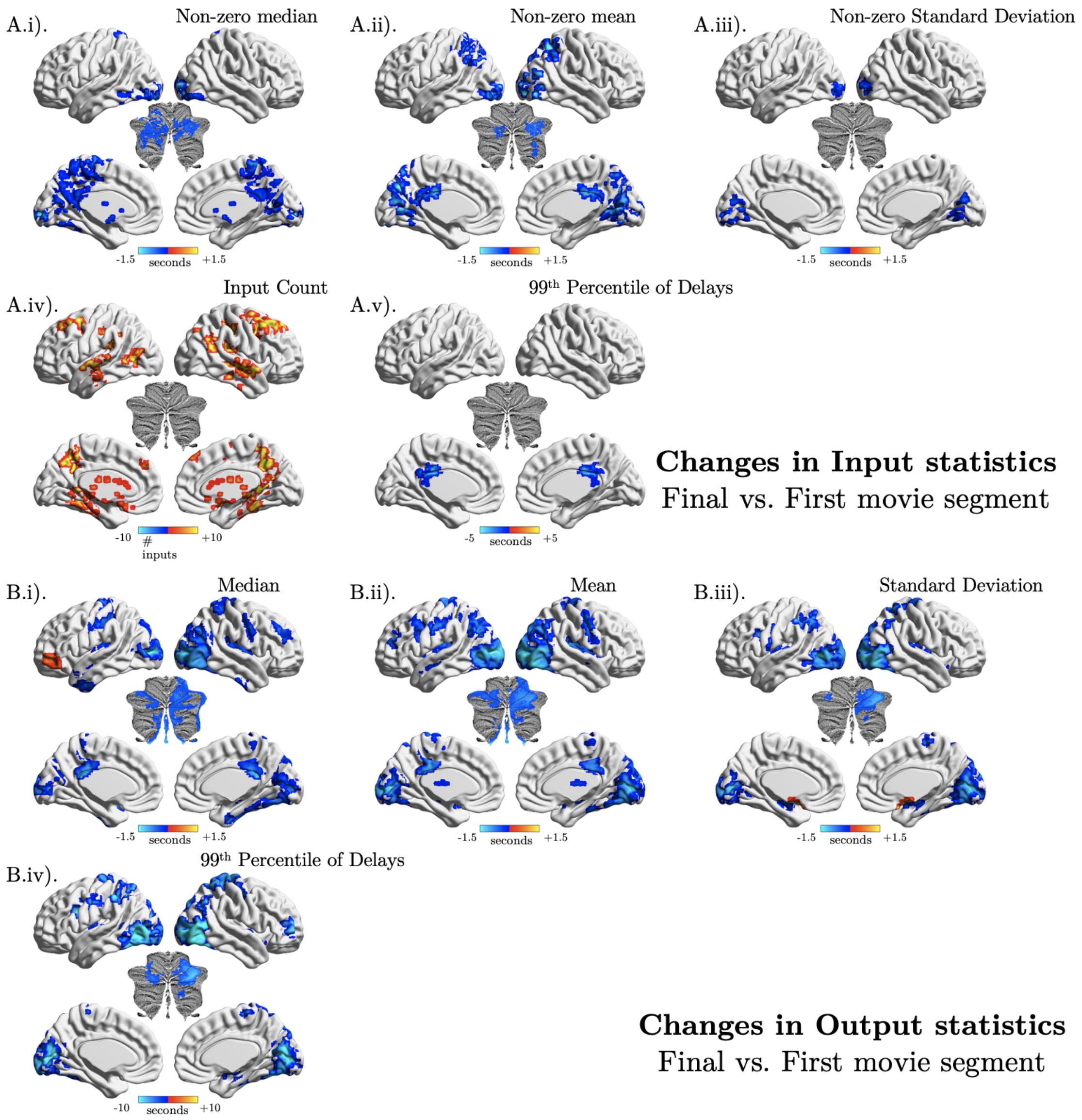


***Supplementary Figure* S2) *Delayed connectivity maps.*** *A group-level linear mixed effects model was used to generate contrasts between the first and final 20 minute segments of 10 movies within A.i) median input delay maps, A.ii) Mean input, A.iii) Standard deviation of inputs, A.iv) count of above-threshold inputs (R ≥ 0.1), A.v) 99th percentile of input delays, B.i), median outputs, B.ii), mean*

*outputs, B.iii, Standard deviation of outputs and B.iv) 99th percentile of output delays. All results are thresholded, corrected for multiple comparisons at alpha = 0.01.*

**Supplementary Table T1)** *Table describing the voxel count and average magnitude of significant clusters revealed through first vs. final 20 minute segment contrasts over input and output connectivity delay maps.*

| Map | Voxel Count | Regions | Average magnitude within cluster |
|---|---|---|---|
| non-zero median inputs (seconds) | 1131 | Posterior Cingulate Cortex, thalamus, paracentral lobule, calcarine gyri | -0.53 (shorter at end of movie) |
| | 520 | LH middle occipital gyrus, LH inferior temporal gyrus, LH lingual gyrus, LH cerebellar crus I | -0.78 |
| | 321 | RH middle occipital gyrus, RH lingual gyrus, RH cerebellar crus I | -1.08 |
| non-zero mean inputs (seconds) | 1648 | Precuneus, calcarine gyri, lingual gyri, middle occipital gyri, LH parietal lobule posterior and middle cingulate cortices | -1.19 |
| | 192 | RH parietal lobule (inferior & superior) | -1.27 |
| | 115 | RH cerebellar crus I & II | -0.91 |
| non-zero standard deviation of inputs (seconds) | 369 | LH & RH middle occipital gyri | -0.53 |
| Number of inputs (count) | 1793 | Precuneus, parieto-occipital sulcus, thalamus, caudate nucleus, LH lingual gyrus, LH fusiform gyus | 7.48 |
| | 1384 | RH middle temporal gyrus, RH superior temporal sulcus, RH rolandic operculum, RH fusiform gyrus | 7.19 |
| | 696 | RH middle frontal & superior medial gyri | 7.94 |
| | 291 | LH middle frontal gyrus | 9.22 |
| | 120 | LH middle temporal gyrus | 11.02 |
| | 115 | LH anterior portion of superior temporal gyrus, LH inferior temporal gyrus | 9.57 |
| 99th percentile of inputs (seconds) | | No differences between first and final movie segments observed | |
| median outputs (seconds) | 3638 | Middle & Superior occipital gyri, RH lingual gyrus, cerebellar crus I, RH cerebellar lobule VIII, | -0.78 |
| | 617 | RH postcentral gyrus, RH Heschl's gyrus, RH posterior insula | -0.45 |

| | | | |
|---|---|---|---|
| | 595 | RH superior parietal lobule | -0.6 |
| | 316 | LH precentral gyrus, LH postcentral gyrus & sulcus | -0.59 |
| | 299 | LH temporal pole, LH insula lobe (posterior), LH thalamus (motor), LH hippocampus | -0.4 |
| | 219 | RH middle frontal gyrus | -0.4 |
| | 177 | RH fusiform gyrus, RH inferior temporal gyrus, RH parahippocampal gyrus | -0.68 |
| | 155 | RH fusiform gyrus, RH inferior temporal gyrus | -0.64 |
| | 134 | LH inferior frontal gyrus (pars orbitalis) | 0.67 |
| mean outputs | 5482 | Middle occipital & calcarine gyri, RH Superior occipital gyrus, LH lingual gyrus, cerebellar crus I, RH cerebellar lobule VIII, | -0.95 |
| | 1337 | LH precentral gyrus, LH postcentral gyrus, LH rolandic operculum, LH pars triangularis, LH insula, LH putamen, LH thalamus | -0.61 |
| | 1209 | RH pre gyrus, RH Heschl's gyrus, RH rolandic operculum, RH posterior insula, RH thalamus | -0.63 |
| | 448 | LH angular gyrus | -0.74 |
| | 309 | Middle cingulate cortex | -0.75 |
| | 195 | RH postcentral gyrus | -0.68 |
| | 113 | left middle frontal gyrus | -0.73 |
| standard deviation of outputs | 3728 | Middle, inferior occipital & calcarine gyri, lingual gyri, RH cerebellar crus I, RH cerebellar lobule VIII | -0.97 |
| | 605 | LH rolandic operculum, inferior portion of LH pre & postcentral gyri & postcentral sulcus, LH hippocampus, LH piriform cortex, LH putamen | -0.73 |
| | 463 | RH heschl's gyrus, RH hippocampus, RH thalamus, RH piriform cortex | -0.56 |
| | 379 | LH superior & inferior parietal lobules | -0.91 |
| | 185 | LH & RH pars orbitalis | 1.0 |
| | 179 | RH postcentral sulcus, RH paracentral lobule | -0.75 |
| | 112 | RH superior parietal lobule | -0.91 |
| 99th percentile of outputs | | Middle occipital & calcarine gyri, lingual gyri, posterior portion of LH inferior temporal gyrus, cerebellar crus I | -4.02 |
| | | Cerebellar lobules VIII & IX | -3.28 |
| | | LH pars opercularis, LH insula lobe | -3.69 |
| | | RH superior parietal lobule | -3.83 |
| | | RH hippocampus, RH parahippocampal gyrus, RH amygdala, RH pallidum | -2.48 |
| | | RH middle frontal gyrus | -4.45 |

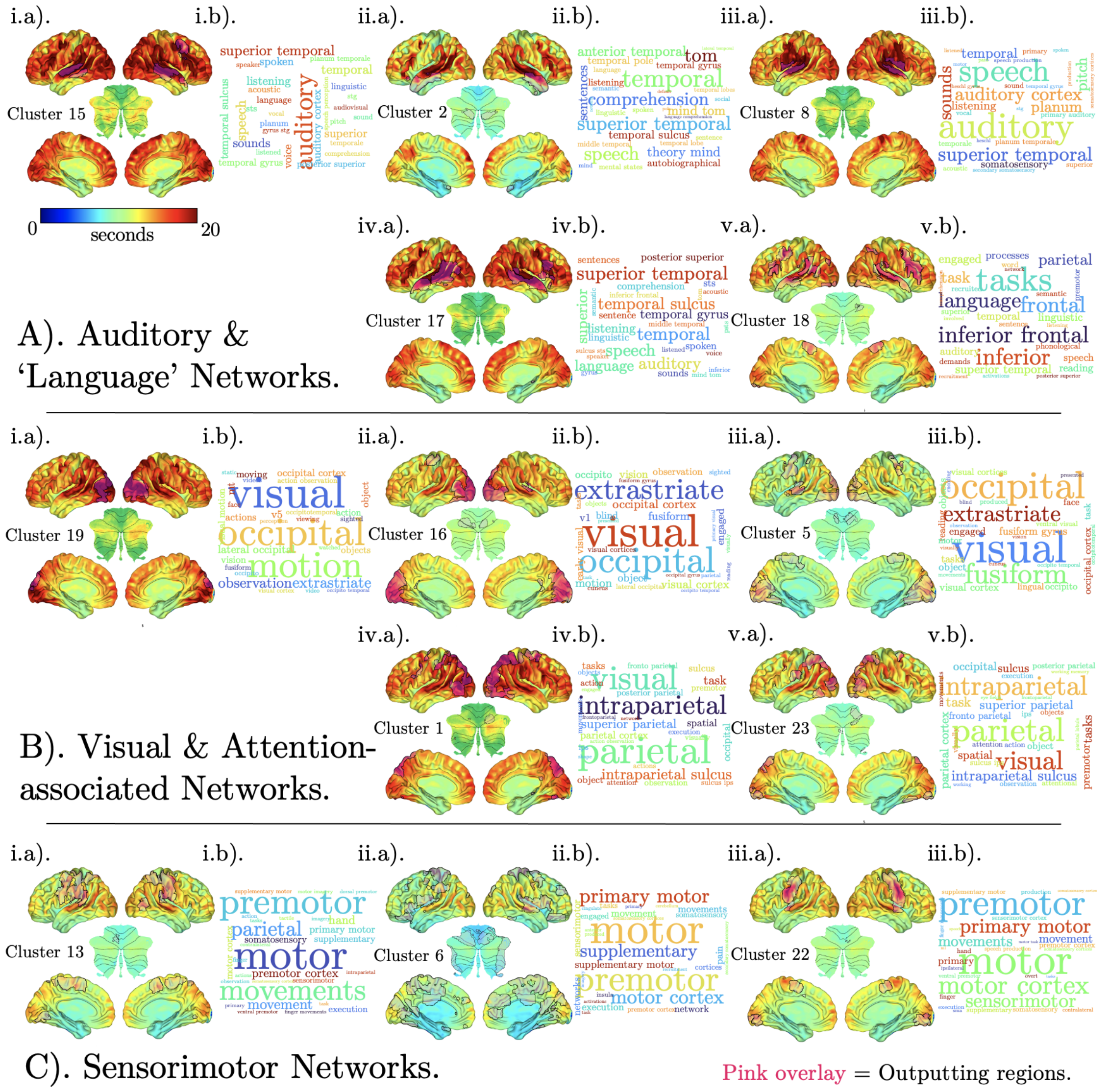


***Supplementary Figure S3) APC clusters, organised by inferred functions of output networks.*** *Each of 807 group-averaged input-delay maps produced from each of 807 unique outputting 'seed' DiFumo ROIs were assigned to one of 24 clusters. These maps were averaged within each cluster, yielding cluster-wise delay maps, overwhich a combined mask of the outputting seed regions were superimposed (displayed in pink over whole-brain surfaces). These cluster + seed maps were organised according to a meta-analysis of terms in the neurosynth database associated with the voxels present in each outputting 'seed' region mask. This yielded 5 categories, the first three of which are*

*presented here. These include A) 'Auditory and Language', comprising 5 cluster maps with overlaid outputting regions (A.i-v.b), presented next to a wordcloud of the top 50 terms yielded from meta-analytic decoding of outputting regions (A.i-v.a)., B) 'Visual and Attention-associated networks', and C) 'Sensorimotor networks'.*

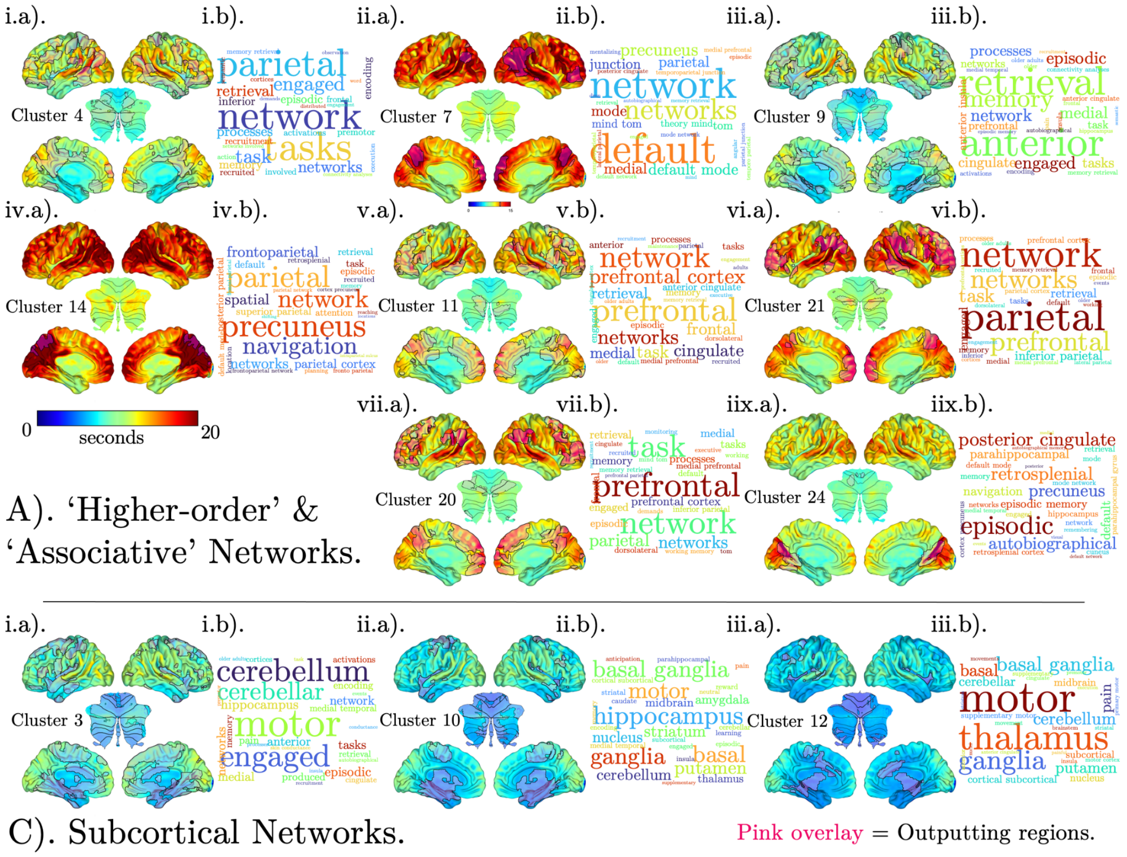


***Supplementary Figure S4)*** *APC clusters continued, organised by meta-analytically inferred functions of output networks, including A). 'Higher-order and Associative Networks' and B) 'Subcortical networks.*

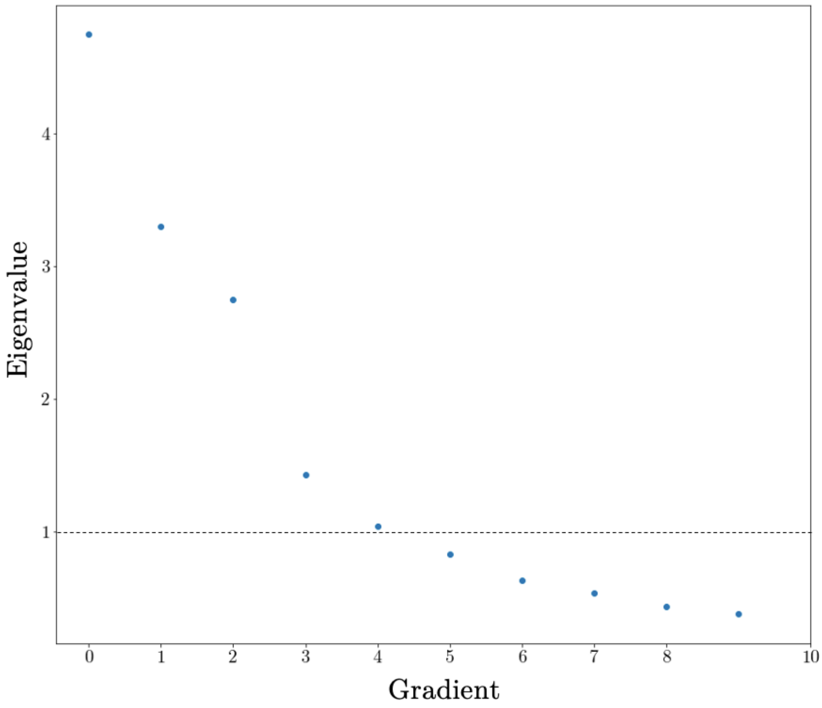


***Supplementary Figure S5)*** *Scree-plot of gradient eigenvalues (λ) against gradients (See Figure 4). A dashed horizontal line is at the threshold of λ = 1. Gradients with λ ≤ 1 were excluded from further consideration.*

***Supplementary Table T5)*** *Global statistics calculated across all voxels within each cluster-wise delay map.*

| | | ***Global Statistics*** | | |
|---|---|---|---|---|
| Cluster | Group | mean | median | std_dev |
| 15 | Auditory & Language Networks | 10.63 | 10.1 | 2.89 |
| 2 | Auditory & Language Networks | 8.31 | 7.72 | 2.62 |
| 8 | Auditory & Language Networks | 9.9 | 9.4 | 2.57 |
| 17 | Auditory & Language Networks | 9.83 | 9.34 | 2.66 |
| 18 | Auditory & Language Networks | 8.79 | 8.48 | 2.03 |
| 19 | Visual & Attention-associated networks | 9.98 | 9.56 | 2.36 |
| 16 | Visual & Attention-associated networks | 8.94 | 8.63 | 1.82 |
| 5 | Visual & Attention-associated networks | 7.49 | 7.21 | 1.51 |
| 1 | Visual & Attention-associated networks | 9.77 | 9.42 | 2.29 |
| 23 | Visual & Attention-associated networks | 8.34 | 8.01 | 1.95 |
| 13 | Sensorimotor Networks | 7.89 | 7.59 | 1.68 |
| 6 | Sensorimotor Networks | 6.95 | 6.61 | 1.58 |
| 22 | Sensorimotor Networks | 8.15 | 7.88 | 1.55 |
| 4 | Higher-order & associative networks | 7.41 | 7.05 | 1.83 |
| 7 | Higher-order & associative networks | 10.1 | 9.61 | 2.65 |
| 9 | Higher-order & associative networks | 6.56 | 6.2 | 1.58 |
| 14 | Higher-order & associative networks | 10.93 | 10.3 | 3.07 |
| 11 | Higher-order & associative networks | 7.91 | 7.54 | 1.97 |
| 21 | Higher-order & associative networks | 9.25 | 8.89 | 2.29 |
| 20 | Higher-order & associative networks | 8.52 | 8.13 | 2.17 |
| 24 | Higher-order & associative networks | 8.01 | 7.61 | 1.92 |
| 3 | Subcortical Networks | 5.96 | 5.63 | 1.34 |
| 10 | Subcortical Networks | 5.37 | 5.06 | 1.11 |
| 12 | Subcortical Networks | 4.86 | 4.58 | 0.89 |

***Supplementary Table T6*) Table of Difumo regions used, organised by the cluster they belong to as revealed by APC.**

| Component | Difumo_names | Yeo_networks17 | Network | Cluster |
|---|---|---|---|---|
| 23 | Fusiform gyrus mid-posterior RH | VisCent | Visual | 1 |
| 25 | Intraparietal sulcus posterior RH | DorsAttnA | Dorsal Attention | 1 |
| 30 | Superior parietal lobule posterior RH | DorsAttnA | Dorsal Attention | 1 |
| 52 | Superior parietal lobule postero-superior RH | DorsAttnA | Dorsal Attention | 1 |
| 173 | Lateral occipital cortex anterior LH | DorsAttnB | Dorsal Attention | 1 |
| 178 | Supero occipital sulcus superior RH | DorsAttnA | Dorsal Attention | 1 |
| 192 | Lateral occipital cortex postero-superior RH | DorsAttnA | Dorsal Attention | 1 |
| 211 | Precuneus postero-superior RH | ContC | Dorsal Attention | 1 |
| 260 | Angular gyrus superior anterior LH | ContB | Frontoparietal | 1 |
| 296 | Parieto-occipital sulcus superior | ContC | Frontoparietal | 1 |
| 329 | Superior parietal lobule superior | DorsAttnB | Dorsal Attention | 1 |
| 338 | Superior occipital sulcus posterior LH | VisCent | Visual | 1 |
| 360 | Lateral occipital sulcus posterior RH | VisCent | Visual | 1 |
| 429 | Intraparietal sulcus superior posterior RH | ContA | Frontoparietal | 1 |
| 473 | Lateral occipital cortex posterior RH | DorsAttnA | Dorsal Attention | 1 |
| 566 | Middle frontal gyrus posterior LH | ContA | Dorsal Attention | 1 |
| 576 | Intraparietal sulcus superior RH | DorsAttnA | Dorsal Attention | 1 |
| 608 | Lateral occipital cortex posterior LH | VisCent | Visual | 1 |
| 612 | Superior occipital gyrus superior | DorsAttnA | Dorsal Attention | 1 |
| 724 | Inferior occipital gyrus anterior RH | VisCent | Visual | 1 |
| 755 | Superior parietal lobule antero-superior RH | DorsAttnB | Dorsal Attention | 1 |
| 797 | Precentral gyrus superior and superior frontal gyrus RH | DorsAttnB | Somatomotor | 1 |
| 799 | Superior parietal lobule antero-inferior RH | DorsAttnB | Dorsal Attention | 1 |
| 810 | Superior parietal lobule postero-superior LH | ContC | Dorsal Attention | 1 |
| 882 | Superior occipital gyrus inferior RH | VisCent | Visual | 1 |
| 887 | Lateral occipital cortex antero-inferior LH | DorsAttnA | Dorsal Attention | 1 |
| 915 | Postcentral gyrus middle posterior LH | DorsAttnB | Dorsal Attention | 1 |
| 923 | Superior occipital sulcus superior LH | DorsAttnA | Dorsal Attention | 1 |

| | | | | |
|---|---|---|---|---|
| 988 | Superior parietal lobule RH | DorsAttnA | Dorsal Attention | 1 |
| 53 | Temporal pole superior RH | DefaultB | Default | 2 |
| 67 | Superior temporal sulcus anterior RH | TempPar | Default | 2 |
| 113 | Middle temporal gyrus mid-anterior lateral LH | DefaultB | Default | 2 |
| 123 | Middle temporal gyrus middle LH | DefaultB | Default | 2 |
| 300 | Cerebellum Crus II | No network found | No network found | 2 |
| 397 | Middle temporal gyrus middle posterior LH | DefaultB | Default | 2 |
| 524 | Temporal pole superior | DefaultB | Limbic | 2 |
| 588 | Superior temporal sulcus mid-anterior RH | TempPar | Default | 2 |
| 9 | Parahippocampal gyrus anterior | No network found | Default | 3 |
| 42 | Cerebellum VI superior | VisPeri | No network found | 3 |
| 48 | Superior frontal sulcus mid-posterior LH | ContB | Default | 3 |
| 54 | Retrosplenial cortex | No network found | Default | 3 |
| 56 | Anterior insula inferior LH | SalVentAttnA | Salience | 3 |
| 65 | Parahippocampal sulcus | DefaultC | Visual | 3 |
| 76 | Fusiform gyrus anterior LH | DefaultC | Visual | 3 |
| 78 | Central sulcus mid-superior LH | SomMotA | Somatomotor | 3 |
| 84 | Cerebellum VIIIab | No network found | No network found | 3 |
| 92 | Hippocampus posterior inferior | DefaultC | Subcortical | 3 |
| 98 | Cerebellum IX middle | No network found | No network found | 3 |
| 106 | Inferior rostral gyrus RH | DefaultA | Default | 3 |
| 112 | Superior frontal gyrus mid-posterior RH | ContB | Frontoparietal | 3 |
| 134 | Subgenual cortex | LimbicB | Limbic | 3 |
| 154 | Central sulcus and postcentral sulcus superior | SomMotA | Somatomotor | 3 |
| 157 | Occipitotemporal sulcus middle RH | DorsAttnA | Visual | 3 |
| 164 | Cerebellum VIIb | No network found | No network found | 3 |
| 166 | Callosal sulcus middle | ContC | Frontoparietal | 3 |
| 175 | Frontomarginal gyrus | DefaultA | Default | 3 |
| 205 | Circular sulcus of the insula posterior LH | SomMotB | Somatomotor | 3 |

| | | | | |
|---|---|---|---|---|
| 209 | Superior frontal sulcus posterior inferior LH | DorsAttnB | Dorsal Attention | 3 |
| 217 | Cerebellum Crus I LH | No network found | No network found | 3 |
| 221 | Planum polare | SomMotB | Salience | 3 |
| 222 | Precentral gyrus mid-superior RH | DorsAttnB | Somatomotor | 3 |
| 234 | Precentral gyrus mid-superior LH | SomMotA | Somatomotor | 3 |
| 235 | Anterior cingulate cortex inferior LH | DefaultA | Default | 3 |
| 239 | Anterior insula anterior RH | SalVentAttnB | Frontoparietal | 3 |
| 250 | Lateral geniculate nucleus LH | No network found | Subcortical | 3 |
| 252 | Cerebellum Crus I LH | No network found | No network found | 3 |
| 264 | Anterior cingulate cortex superior | SalVentAttnB | Salience | 3 |
| 266 | Parieto-occipital sulcus anterior | DefaultC | Visual | 3 |
| 277 | Superior frontal gyrus middle LH | DefaultA | Default | 3 |
| 286 | Superior frontal sulcus mid-anterior RH | DefaultA | Default | 3 |
| 299 | Middle frontal gyrus mid-posterior superior RH | ContB | Frontoparietal | 3 |
| 301 | Superior frontal gyrus middle | SalVentAttnB | Salience | 3 |
| 309 | Precentral gyrus inferior | SomMotB | Somatomotor | 3 |
| 347 | Cerebellum Crus I RH | No network found | No network found | 3 |
| 355 | Frontomarginal sulcus RH | ContB | Frontoparietal | 3 |
| 362 | Lingual gyrus middle LH | VisPeri | Visual | 3 |
| 363 | Inferior frontal gyrus mid-posterior LH | ContA | Frontoparietal | 3 |
| 372 | Cuneus anterior RH | VisPeri | Visual | 3 |
| 394 | Dorsal visual stream posterior RH | DorsAttnA | Dorsal Attention | 3 |
| 398 | Cingulate sulcus posterior | SalVentAttnA | Salience | 3 |
| 407 | Posterior cingulate cortex inferior LH | DefaultC | Default | 3 |
| 409 | Cerebellum IX | No network found | No network found | 3 |
| 411 | Cerebellum Vermis IX | No network found | No network found | 3 |
| 431 | Paracentral lobule inferior LH | SomMotA | Somatomotor | 3 |
| 447 | Collateral sulcus anterior | LimbicA | Limbic | 3 |
| 449 | Calcarine sulcus anterior LH | VisPeri | Visual | 3 |

| | | | | |
|---|---|---|---|---|
| 457 | Posterior insula RH | SomMotB | Somatomotor | 3 |
| 500 | Planum polare LH | SomMotB | Somatomotor | 3 |
| 517 | Superior frontal gyrus posterior medial RH | SalVentAttnA | Salience | 3 |
| 543 | Superior temporal gyrus anterior LH | LimbicA | Limbic | 3 |
| 544 | Postcentral sulcus mid-superior LH | DorsAttnB | Dorsal Attention | 3 |
| 558 | Heschl's gyrus medial RH | SomMotB | Somatomotor | 3 |
| 597 | Posterior insula inferior RH | SalVentAttnA | Salience | 3 |
| 625 | Cerebellum VIIIa LH | No network found | No network found | 3 |
| 648 | Superior temporal gyrus anterior medial RH | LimbicA | Limbic | 3 |
| 668 | Occipitotemporal sulcus middle LH | No network found | Visual | 3 |
| 682 | Optic chiasm anterior | LimbicB | Limbic | 3 |
| 696 | Superior frontal gyrus posterior superior LH | SalVentAttnA | Salience | 3 |
| 699 | Superior frontal gyrus anterior | DefaultB | Default | 3 |
| 708 | Calcarine sulcus mid-anterior RH | VisPeri | Visual | 3 |
| 717 | Subcallosal cortex superior | LimbicB | Limbic | 3 |
| 751 | Lateral fissure middle LH | SomMotB | Somatomotor | 3 |
| 753 | Precentral sulcus medial RH | SomMotA | Somatomotor | 3 |
| 754 | Lingual gyrus anterior LH | VisPeri | Visual | 3 |
| 759 | Subcallosal cortex inferior | LimbicB | Limbic | 3 |
| 774 | Calcarine sulcus anterior RH | VisPeri | Visual | 3 |
| 784 | Cerebellum Crus I LH | No network found | No network found | 3 |
| 793 | Posterior insula inferior LH | SalVentAttnA | Salience | 3 |
| 795 | Occipitotemporal sulcus anterior RH | LimbicA | Visual | 3 |
| 804 | Lateral orbital gyrus RH | LimbicB | Limbic | 3 |
| 815 | Cerebellum IX | No network found | No network found | 3 |
| 820 | Suborbital sulcus LH | LimbicB | Limbic | 3 |
| 822 | Lingual gyrus anterior RH | VisPeri | Visual | 3 |
| 827 | Temporal pole inferior | LimbicA | Limbic | 3 |
| 841 | Posterior orbital gyrus RH | LimbicB | Limbic | 3 |

| | | | | |
|---|---|---|---|---|
| 848 | Paracingulate sulcus superior RH | SalVentAttnB | Salience | 3 |
| 855 | Inferior temporal sulcus mid-anterior LH | DefaultB | Default | 3 |
| 889 | Cerebellum IX | No network found | No network found | 3 |
| 891 | Cerebellum V RH | No network found | No network found | 3 |
| 910 | Cerebellum horizontal fissure | No network found | No network found | 3 |
| 920 | Cerebellum VIIb | No network found | No network found | 3 |
| 924 | Inferior frontal gyrus mid-anterior RH | SalVentAttnB | Frontoparietal | 3 |
| 930 | Cerebellum VI LH | No network found | No network found | 3 |
| 933 | Cerebellum V RH | DorsAttnA | No network found | 3 |
| 937 | Collateral sulcus posterior RH | VisCent | Visual | 3 |
| 962 | Paracentral lobule postero-inferior | SomMotA | Somatomotor | 3 |
| 965 | Paracentral lobule posterior LH | SomMotA | Somatomotor | 3 |
| 966 | Paracingulate sulcus posterior LH | SalVentAttnA | Salience | 3 |
| 981 | Posterior insula postero-superior RH | SomMotB | Somatomotor | 3 |
| 985 | Cerebellum Crus I RH | No network found | No network found | 3 |
| 990 | Caudate posterior RH | No network found | Subcortical | 3 |
| 1001 | Cerebellum VI RH | No network found | No network found | 3 |
| 1003 | Cerebellum VI | No network found | No network found | 3 |
| 1023 | Caudate superior anterior | No network found | Subcortical | 3 |
| 62 | Ventromedial prefrontal cortex posterior | LimbicB | Limbic | 4 |
| 70 | Posterior cingulate cortex anterior | DefaultA | Default | 4 |
| 97 | Lateral occipital cortex mid-anterior RH | DefaultC | Dorsal Attention | 4 |
| 103 | Inferior temporal sulcus mid-posterior LH | ContB | Frontoparietal | 4 |
| 104 | Cingulate sulcus mid-posterior | SalVentAttnA | Salience | 4 |
| 114 | Callosomarginal sulcus inferior | SomMotA | Somatomotor | 4 |
| 130 | Collateral sulcus posterior | DefaultC | Visual | 4 |
| 135 | Inferior frontal gyrus anterior RH | ContB | Frontoparietal | 4 |

| | | | | |
|---|---|---|---|---|
| 179 | Occipitotemporal sulcus mid-posterior LH | DorsAttnA | Dorsal Attention | 4 |
| 226 | Lateral occipital cortex superior LH | DefaultC | Dorsal Attention | 4 |
| 229 | Frontal opercula medial | SalVentAttnA | Salience | 4 |
| 230 | Anterior occipital sulcus superior RH | DorsAttnA | Dorsal Attention | 4 |
| 245 | Fusiform gyrus mid-anterior LH | VisCent | Visual | 4 |
| 259 | Middle temporal gyrus postero-superior RH | TempPar | Salience | 4 |
| 279 | Supramarginal gyrus antero-inferior LH | SalVentAttnA | Salience | 4 |
| 346 | Frontal pole medial LH | DefaultB | Default | 4 |
| 356 | Orbital H-shaped sulcus RH | ContB | Frontoparietal | 4 |
| 368 | Superior occipital gyrus superior LH | DorsAttnA | Visual | 4 |
| 402 | Anterior occipital sulcus anterior RH | DorsAttnA | Dorsal Attention | 4 |
| 481 | Precentral gyrus middle anterior RH | SalVentAttnA | Salience | 4 |
| 490 | Middle temporal gyrus mid-posterior medial LH | DefaultB | Default | 4 |
| 499 | Inferior temporal gyrus posterior RH | DorsAttnA | Dorsal Attention | 4 |
| 550 | Postcentral gyrus middle LH | SomMotA | Somatomotor | 4 |
| 552 | Cerebellum Crus I LH | No network found | No network found | 4 |
| 564 | Cerebellum VI | No network found | No network found | 4 |
| 590 | Subparietal sulcus anterior RH | ContC | Frontoparietal | 4 |
| 618 | Superior parietal sulcus superior LH | DorsAttnB | Dorsal Attention | 4 |
| 634 | Precuneus inferior LH | DefaultA | Default | 4 |
| 641 | Cerebellum Crus II | No network found | No network found | 4 |
| 649 | Cerebellum Crus II LH | No network found | No network found | 4 |
| 657 | Middle frontal gyrus middle inferior LH | ContA | Frontoparietal | 4 |
| 660 | Postcentral gyrus superior LH | SomMotA | Somatomotor | 4 |
| 710 | Posterior orbital gyrus posterior | DefaultB | Default | 4 |
| 713 | Superior temporal sulcus postero-superior LH | TempPar | Salience | 4 |
| 716 | Intraparietal sulcus mid-posterior LH | ContA | Frontoparietal | 4 |
| 720 | Middle temporal gyrus anterior RH | DefaultA | Default | 4 |
| 745 | Subparietal sulcus anterior LH | ContC | Default | 4 |

| | | | | |
|---|---|---|---|---|
| 766 | Precentral gyrus inferior RH | DorsAttnB | Dorsal Attention | 4 |
| 773 | Inferior frontal sulcus middle RH | ContA | Frontoparietal | 4 |
| 786 | Inferior temporal sulcus posterior RH | ContB | Frontoparietal | 4 |
| 803 | Superior frontal sulcus posterior LH | ContA | Frontoparietal | 4 |
| 811 | Superior parietal sulcus inferior LH | ContA | Dorsal Attention | 4 |
| 834 | Middle frontal sulcus mid-anterior LH | SalVentAttnB | Frontoparietal | 4 |
| 838 | Posterior cingulate cortex antero-inferior | ContC | Default | 4 |
| 847 | Postcentral gyrus superior RH | SomMotA | Somatomotor | 4 |
| 860 | Precuneus antero-superior LH | SalVentAttnA | Salience | 4 |
| 873 | Cerebellum VI RH | VisCent | Visual | 4 |
| 901 | Cerebellum Crus II LH | No network found | No network found | 4 |
| 902 | Cingulate gyrus mid-posterior | ContC | Default | 4 |
| 943 | Inferior frontal sulcus posterior superior LH | ContB | Frontoparietal | 4 |
| 946 | Inferior frontal sulcus posterior inferior LH | ContA | Frontoparietal | 4 |
| 976 | Lateral orbital gyrus LH | DefaultB | Default | 4 |
| 1002 | Middle frontal gyrus posterior superior LH | ContB | Frontoparietal | 4 |
| 1 | Retrocalcarine cortex RH | VisCent | Visual | 5 |
| 33 | Parieto-occipital sulcus middle LH | VisPeri | Visual | 5 |
| 60 | Calcarine sulcus posterior LH | VisCent | Visual | 5 |
| 71 | Heschl's gyrus posterior LH | SomMotB | Somatomotor | 5 |
| 75 | Calcarine cortex anterior RH | VisPeri | Visual | 5 |
| 117 | Lingual gyrus mid-posterior RH | VisPeri | Visual | 5 |
| 177 | Inferior occipital gyrus LH | VisCent | Visual | 5 |
| 197 | Cuneus anterior LH | VisPeri | Visual | 5 |
| 233 | Precentral gyrus antero-superior LH | SomMotA | Somatomotor | 5 |
| 243 | Parieto-occipital sulcus mid-anterior LH | DefaultC | Visual | 5 |
| 265 | Precentral sulcus mid-superior RH | DorsAttnB | Dorsal Attention | 5 |
| 271 | Inferior occipital sulcus superior RH | VisCent | Visual | 5 |
| 288 | Descending occipital gyrus superior | VisPeri | Visual | 5 |
| 297 | Descending occipital gyrus superior LH | VisCent | Visual | 5 |

| | | | | |
|---|---|---|---|---|
| 306 | Intracalcarine cortex LH | VisPeri | Visual | 5 |
| 391 | Collateral sulcus posterior LH | VisCent | Visual | 5 |
| 401 | Descending occipital gyrus inferior LH | VisCent | Visual | 5 |
| 436 | Precentral gyrus middle posterior LH | SomMotB | Somatomotor | 5 |
| 466 | Calcarine cortex mid-anterior LH | VisPeri | Visual | 5 |
| 508 | Paracentral lobule superior | SomMotA | Somatomotor | 5 |
| 549 | Collateral sulcus middle RH | VisCent | Visual | 5 |
| 605 | Cuneus inferior RH | VisPeri | Visual | 5 |
| 662 | Occipitotemporal gyrus posterior LH | DorsAttnA | Visual | 5 |
| 677 | Lingual gyrus middle RH | VisPeri | Visual | 5 |
| 692 | Fusiform gyrus middle RH | VisCent | Visual | 5 |
| 761 | Fusiform gyrus RH | VisPeri | Visual | 5 |
| 777 | Superior occipital sulcus inferior RH | VisCent | Visual | 5 |
| 780 | Lingual gyrus middle | VisPeri | Visual | 5 |
| 833 | Calcarine sulcus mid-anterior | VisPeri | Visual | 5 |
| 898 | Collateral sulcus mid-posterior LH | VisCent | Visual | 5 |
| 932 | Occipitotemporal gyrus posterior RH | DorsAttnA | Visual | 5 |
| 952 | Planum temporale LH | SomMotB | Somatomotor | 5 |
| 49 | Cerebellum VI superior RH | VisPeri | Visual | 6 |
| 68 | Dorsomedial prefrontal cortex RH | DefaultB | Default | 6 |
| 100 | Central sulcus inferior | SomMotB | Somatomotor | 6 |
| 124 | Callosomarginal sulcus superior RH | DorsAttnB | Somatomotor | 6 |
| 127 | Superior parts of central and postcentral sulci LH | SomMotA | Somatomotor | 6 |
| 133 | Occipitotemporal sulcus mid-anterior RH | DorsAttnA | Dorsal Attention | 6 |
| 161 | Cingulate cortex mid-posterior | DefaultA | Salience | 6 |
| 202 | Subcentral gyrus LH | SomMotB | Somatomotor | 6 |
| 212 | Cingulate anterior RH | DefaultA | Default | 6 |
| 225 | Parietal operculum anterior RH | SomMotB | Somatomotor | 6 |
| 244 | Paracentral lobule inferior | SomMotA | Somatomotor | 6 |
| 257 | Heschl's gyrus middle RH | SomMotB | Somatomotor | 6 |
| 304 | Supramarginal gyrus antero-superior LH | DorsAttnB | Dorsal Attention | 6 |

| | | | | |
|---|---|---|---|---|
| 316 | Lateral fissure posterior | SomMotB | Somatomotor | 6 |
| 322 | Insula middle superior | SalVentAttnA | Salience | 6 |
| 326 | Precentral gyrus superior | SomMotA | Somatomotor | 6 |
| 334 | Pars triangularis LH | DefaultB | Default | 6 |
| 335 | Posterior orbital gyrus | LimbicB | Limbic | 6 |
| 352 | Cerebellum VI LH | VisCent | No network found | 6 |
| 389 | Superior frontal gyrus anterior RH | SalVentAttnB | Default | 6 |
| 415 | Pars opercularis RH | SalVentAttnB | Frontoparietal | 6 |
| 477 | Cingulate middle anterior RH | SalVentAttnB | Frontoparietal | 6 |
| 482 | Operculum orbitalis LH | SalVentAttnB | Salience | 6 |
| 520 | Cingulate posterior | SomMotA | Somatomotor | 6 |
| 521 | Cuneus superior LH | VisPeri | Visual | 6 |
| 529 | Retrosplenial cortex posterior | DefaultC | Default | 6 |
| 592 | Central operculum superior LH | SomMotB | Somatomotor | 6 |
| 639 | Cerebellum Crus I anterior RH | No network found | No network found | 6 |
| 651 | Superior parietal lobule antero-superior LH | DorsAttnB | Dorsal Attention | 6 |
| 671 | Precuneus antero-superior | ContC | Dorsal Attention | 6 |
| 680 | Parietal operculum medial RH | SomMotB | Somatomotor | 6 |
| 693 | Lateral orbital gyrus anterior | DefaultB | Default | 6 |
| 697 | Central sulcus mid-superior RH | SomMotA | Somatomotor | 6 |
| 721 | Postcentral gyrus superior | SomMotA | Somatomotor | 6 |
| 723 | Anterior orbital gyrus lateral | LimbicB | Frontoparietal | 6 |
| 727 | Central sulcus middle medial RH | SomMotA | Somatomotor | 6 |
| 731 | Cingulate mid-posterior | SalVentAttnA | Somatomotor | 6 |
| 736 | Central sulcus superior LH | SomMotA | Somatomotor | 6 |
| 737 | Interhemispheric fissure precentral gyrus | SalVentAttnA | Salience | 6 |
| 756 | Postcentral sulcus superior LH | SomMotA | Somatomotor | 6 |
| 767 | Superior parts of central and postcentral gyrus | SomMotA | Somatomotor | 6 |
| 768 | Calcarine sulcus middle | VisPeri | Visual | 6 |
| 783 | Precentral gyrus superior RH | SomMotA | Somatomotor | 6 |

| | | | | |
|---|---|---|---|---|
| 845 | Precentral sulcus inferior RH | ContA | Salience | 6 |
| 867 | Subparietal sulcus inferior RH | DefaultC | Default | 6 |
| 883 | Superior frontal sulcus mid-anterior LH | DefaultA | Default | 6 |
| 896 | Lingual gyrus medial | VisPeri | Visual | 6 |
| 904 | Parieto-occipital sulcus posterior RH | VisPeri | Visual | 6 |
| 960 | Central sulcus superior RH | SomMotA | Somatomotor | 6 |
| 968 | Callosal sulcus mid-posterior | ContC | Frontoparietal | 6 |
| 970 | Callosomarginal sulcus superior | SomMotA | Somatomotor | 6 |
| 974 | Middle frontal gyrus posterior superior RH | ContA | Frontoparietal | 6 |
| 992 | Postcentral sulcus medial | SomMotA | Somatomotor | 6 |
| 996 | Cerebellum VI LH | No network found | No network found | 6 |
| 101 | Lateral occipital cortex superior RH | ContB | Frontoparietal | 7 |
| 128 | Precuneus middle LH | ContC | Default | 7 |
| 215 | Frontal pole medial | DefaultA | Default | 7 |
| 231 | Angular sulcus mid-anterior RH | DefaultA | Default | 7 |
| 332 | Angular gyrus posterior inferior RH | DefaultC | Default | 7 |
| 396 | Superior frontal gyrus anterior medial | DefaultB | Default | 7 |
| 433 | Angular gyrus postero-inferior RH | TempPar | Dorsal Attention | 7 |
| 446 | Angular gyrus postero-superior RH | ContB | Frontoparietal | 7 |
| 561 | Precuneus middle superior LH | ContC | Frontoparietal | 7 |
| 568 | Orbital cortex lateral RH | ContB | Frontoparietal | 7 |
| 676 | Precuneus posterior | ContC | Default | 7 |
| 738 | Angular gyrus posterior RH | DefaultA | Default | 7 |
| 762 | Angular sulcus posterior RH | DefaultA | Default | 7 |
| 764 | Angular gyrus antero-superior RH | ContB | Frontoparietal | 7 |
| 832 | Precuneus posterior RH | ContC | Frontoparietal | 7 |
| 850 | Angular sulcus mid-posterior RH | DefaultC | Dorsal Attention | 7 |
| 928 | Angular sulcus posterior LH | DefaultA | Default | 7 |
| 989 | Superior occipital gyrus RH | DefaultC | Default | 7 |
| 16 | Subcentral gyrus RH | SomMotB | Somatomotor | 8 |

| | | | | |
|---|---|---|---|---|
| 77 | Supramarginal gyrus antero-inferior RH | SalVentAttnA | Salience | 8 |
| 83 | Planum temporale anterior LH | SomMotB | Somatomotor | 8 |
| 416 | Heschl's gyrus anterior LH | SomMotB | Somatomotor | 8 |
| 487 | Superior temporal gyrus middle superior LH | SomMotB | Somatomotor | 8 |
| 8 | Inferior temporal sulcus anterior RH | DefaultB | Limbic | 9 |
| 15 | Postcentral sulcus superior RH | DorsAttnB | Somatomotor | 9 |
| 26 | Anterior insula antero-inferior RH | SalVentAttnB | Salience | 9 |
| 36 | Cingulate cortex mid-anterior LH | SalVentAttnB | Salience | 9 |
| 39 | Fusiform gyrus anterior RH | VisCent | Visual | 9 |
| 41 | Central opercular cortex LH | SomMotB | Salience | 9 |
| 111 | Hippocampus posterior superior | No network found | Subcortical | 9 |
| 152 | Middle temporal gyrus mid-anterior medial LH | DefaultB | Default | 9 |
| 156 | Cingulate sulcus mid-anterior LH | SalVentAttnA | Salience | 9 |
| 169 | Temporal pole LH | LimbicA | Limbic | 9 |
| 171 | Anterior insula LH | SalVentAttnA | Salience | 9 |
| 199 | Dorsal visual stream posterior LH | ContA | Dorsal Attention | 9 |
| 204 | Cingulate cortex mid-anterior | SalVentAttnA | Salience | 9 |
| 223 | Subparietal sulcus RH | DefaultA | Default | 9 |
| 246 | Inferior temporal sulcus middle RH | DefaultB | Default | 9 |
| 263 | Cingulate middle posterior RH | SalVentAttnA | Salience | 9 |
| 273 | Anterior insula anterior LH | DefaultB | Default | 9 |
| 291 | Cerebellum VIIb | No network found | No network found | 9 |
| 303 | Pars triangularis superior RH | SalVentAttnB | Frontoparietal | 9 |
| 305 | Posterior insula superior LH | SalVentAttnA | Salience | 9 |
| 308 | Heschl's gyrus anterior medial RH | SomMotB | Somatomotor | 9 |
| 349 | Callosomarginal sulcus middle LH | SalVentAttnA | Salience | 9 |
| 393 | Anterior cingulate cortex inferior | DefaultA | Default | 9 |
| 413 | Frontal operculum RH | SalVentAttnB | Salience | 9 |
| 414 | Anterior orbital gyrus LH | ContB | Frontoparietal | 9 |
| 424 | Pars opercularis superior LH | ContA | Frontoparietal | 9 |

| | | | | |
|---|---|---|---|---|
| 425 | Amygdala anterior | No network found | Subcortical | 9 |
| 452 | Paracingulate gyrus anterior LH | DefaultA | Default | 9 |
| 456 | Cerebellum VI LH | No network found | No network found | 9 |
| 471 | Lingual gyrus mid-anterior LH | VisPeri | Visual | 9 |
| 472 | Frontomarginal sulcus LH | ContB | Frontoparietal | 9 |
| 475 | Paracingulate gyrus mid-anterior LH | DefaultA | Default | 9 |
| 484 | Middle frontal gyrus mid-anterior LH | ContA | Frontoparietal | 9 |
| 502 | Cerebellum Crus II RH | No network found | No network found | 9 |
| 512 | Hippocampal gyrus posterior LH | DefaultC | Visual | 9 |
| 513 | Posterior insula inferior | SalVentAttnA | Salience | 9 |
| 531 | Cerebellum Crus I RH | No network found | No network found | 9 |
| 551 | Lateral fissure anterior RH | SalVentAttnB | Salience | 9 |
| 567 | Anterior insula antero-superior | SalVentAttnB | Salience | 9 |
| 573 | Superior occipital gyrus inferior LH | DorsAttnA | Visual | 9 |
| 578 | Central opercula | SalVentAttnA | Salience | 9 |
| 581 | Superior frontal gyrus middle RH | DefaultB | Default | 9 |
| 584 | Middle temporal gyrus mid-anterior RH | ContB | Frontoparietal | 9 |
| 591 | Occipitotemporal sulcus posterior RH | VisCent | Visual | 9 |
| 604 | Hippocampal gyrus | DefaultC | Visual | 9 |
| 623 | Supramarginal gyrus RH | SalVentAttnB | Frontoparietal | 9 |
| 631 | Superior frontal gyrus antero-superior | DefaultB | Default | 9 |
| 673 | Calcarine sulcus mid-posterior LH | No network found | Visual | 9 |
| 675 | Dorsolateral prefrontal cortex LH | DefaultB | Default | 9 |
| 689 | Middle temporal gyrus mid-anterior LH | DefaultB | Default | 9 |
| 694 | Superior frontal gyrus mid-posterior LH | DefaultB | Default | 9 |
| 707 | Cingulate middle RH | SalVentAttnB | Salience | 9 |
| 711 | Dorsal visual stream RH | ContA | Dorsal Attention | 9 |
| 734 | Middle frontal gyrus mid-posterior LH | ContB | Default | 9 |
| 743 | Subparietal sulcus posterior | DefaultA | Default | 9 |

| | | | | |
|---|---|---|---|---|
| 749 | Callosomarginal sulcus inferior LH | SalVentAttnA | Salience | 9 |
| 757 | Occipitotemporal sulcus posterior LH | DorsAttnA | Dorsal Attention | 9 |
| 775 | Callosomarginal sulcus inferior RH | SalVentAttnA | Salience | 9 |
| 790 | Lateral fissure anterior LH | SomMotB | Salience | 9 |
| 825 | Cerebellum Crus I anterior LH | DorsAttnA | No network found | 9 |
| 826 | Parieto-occipital sulcus postero-superior RH | ContC | Dorsal Attention | 9 |
| 830 | Heschl's gyrus inferior LH | SomMotB | Somatomotor | 9 |
| 840 | Anterior orbital gyrus medial | LimbicB | Limbic | 9 |
| 856 | Superior frontal sulcus anterior LH | DefaultA | Default | 9 |
| 859 | Circular sulcus of the insula antero-superior LH | SalVentAttnA | Salience | 9 |
| 874 | Hippocampus middle RH | No network found | Subcortical | 9 |
| 879 | Middle frontal sulcus middle RH | SalVentAttnB | Salience | 9 |
| 906 | Superior frontal gyrus medial mid-anterior LH | DefaultB | Frontoparietal | 9 |
| 913 | Lingual gyrus anterior | VisPeri | Visual | 9 |
| 914 | Cuneus antero-superior RH | VisPeri | Visual | 9 |
| 921 | Anterior corona radiata anterior RH | SalVentAttnB | Salience | 9 |
| 973 | Suborbital sulcus | DefaultA | Default | 9 |
| 983 | Inferior temporal sulcus medial LH | DorsAttnA | Dorsal Attention | 9 |
| 986 | Superior parts of central and precentral sulci RH | SomMotA | Somatomotor | 9 |
| 1006 | Temporal pole RH | LimbicA | Limbic | 9 |
| 40 | Precentral sulcus middle LH | DorsAttnB | Dorsal Attention | 10 |
| 50 | Thalamus inferior | No network found | Subcortical | 10 |
| 58 | Hippocampal fissure | DefaultC | Subcortical | 10 |
| 64 | Cerebellum V LH | No network found | No network found | 10 |
| 69 | Putamen superior RH | No network found | Subcortical | 10 |
| 82 | Uncinate fasciculus | No network found | Limbic | 10 |
| 95 | Thalamus superior | No network found | Subcortical | 10 |
| 147 | Caudate posterior LH | No network found | Subcortical | 10 |

| | | | | |
|---|---|---|---|---|
| 148 | Circular sulcus of the insula mid-posterior RH | SomMotB | Somatomotor | 10 |
| 149 | Putamen anterior RH | No network found | Subcortical | 10 |
| 153 | Caudate | No network found | Subcortical | 10 |
| 170 | Cerebellum VI | No network found | No network found | 10 |
| 181 | Globus pallidus inferior | No network found | Subcortical | 10 |
| 191 | Superior frontal gyrus posterior RH | DorsAttnB | Dorsal Attention | 10 |
| 201 | Superior temporal gyrus medial RH | LimbicA | Limbic | 10 |
| 216 | Cerebellum X | No network found | No network found | 10 |
| 256 | Fornix superior | No network found | Subcortical | 10 |
| 261 | Cerebellum Vermis X | No network found | No network found | 10 |
| 280 | Midbrain anterior | No network found | Subcortical | 10 |
| 283 | Cerebellum VI | No network found | No network found | 10 |
| 287 | Collateral sulcus anterior LH | DefaultC | Limbic | 10 |
| 302 | Cerebellum IV | DefaultC | Visual | 10 |
| 317 | Optic chiasm | No network found | Subcortical | 10 |
| 319 | Cerebellum VIIb | No network found | No network found | 10 |
| 327 | Precentral sulcus medial inferior LH | SomMotA | Somatomotor | 10 |
| 333 | Hippocampal gyrus RH | LimbicA | Limbic | 10 |
| 337 | Superior frontal sulcus posterior RH | DorsAttnB | Dorsal Attention | 10 |
| 358 | Cerebellum VIIIa | No network found | No network found | 10 |
| 370 | Entorhinal cortex | DefaultC | Limbic | 10 |
| 379 | Central sulcus of the insula postero-inferior | TempPar | Somatomotor | 10 |
| 386 | Cingulum inferior | No network found | Subcortical | 10 |
| 395 | Middle frontal sulcus posterior RH | ContA | Frontoparietal | 10 |
| 420 | Cerebellum Vermis | VisPeri | No network found | 10 |
| 426 | Caudate LH | No network found | Subcortical | 10 |

| | | | | |
|---|---|---|---|---|
| 460 | Inferior temporal gyrus anterior RH | LimbicA | Limbic | 10 |
| 476 | Cerebellum IV | No network found | No network found | 10 |
| 480 | Parahippocampal gyrus posterior RH | VisPeri | Default | 10 |
| 485 | Caudate superior posterior | No network found | Subcortical | 10 |
| 486 | Corpus callosum genu inferior | No network found | Subcortical | 10 |
| 488 | Globus pallidus middle | No network found | Subcortical | 10 |
| 504 | Temporal pole superior medial LH | LimbicA | Limbic | 10 |
| 505 | Precentral gyrus medial | SomMotA | Somatomotor | 10 |
| 530 | Circular sulcus of the insula inferior | No network found | Somatomotor | 10 |
| 559 | Insula inferior RH | SalVentAttnA | Salience | 10 |
| 607 | Occipitotemporal gyrus posterior | No network found | Visual | 10 |
| 614 | Superior temporal gyrus anterior medial | SalVentAttnA | Salience | 10 |
| 615 | Caudate anterior RH | No network found | Subcortical | 10 |
| 635 | Cerebellum VI RH | No network found | No network found | 10 |
| 644 | Putamen posterior LH | SalVentAttnA | Salience | 10 |
| 653 | Collateral sulcus middle LH | DefaultC | Limbic | 10 |
| 655 | Precentral sulcus medial LH | SomMotA | Somatomotor | 10 |
| 678 | Amygdala posterior | No network found | Subcortical | 10 |
| 725 | Anterior insula mid-inferior LH | SalVentAttnB | Salience | 10 |
| 726 | Parahippocampal gyrus middle RH | DefaultC | Limbic | 10 |
| 750 | Superior parietal sulcus superior RH | DorsAttnB | Dorsal Attention | 10 |
| 779 | Circular sulcus of the insula posterior RH | No network found | Somatomotor | 10 |
| 798 | Ventral striatum RH | LimbicB | Limbic | 10 |
| 823 | Occipitotemporal gyrus anterior LH | LimbicA | Limbic | 10 |
| 836 | Cerebellum Crus I | No network found | No network found | 10 |
| 842 | Precentral gyrus superior LH | SomMotA | Somatomotor | 10 |

| | | | | |
|---|---|---|---|---|
| 851 | Putamen anterior LH | No network found | Subcortical | 10 |
| 877 | Hippocampus anterior LH | No network found | Subcortical | 10 |
| 890 | Putamen anterior | No network found | Subcortical | 10 |
| 907 | Paracentral sulcus superior | SomMotA | Somatomotor | 10 |
| 912 | Cerebellum VI RH | No network found | No network found | 10 |
| 919 | Cingulate sulcus posterior RH | SomMotA | Somatomotor | 10 |
| 987 | Cerebellum VI RH | No network found | No network found | 10 |
| 63 | Superior frontal sulcus middle LH | ContB | Default | 11 |
| 86 | Cingulate mid-anterior RH | DefaultA | Default | 11 |
| 90 | Middle frontal gyrus middle LH | SalVentAttnB | Salience | 11 |
| 116 | Pars opercularis LH | DefaultB | Default | 11 |
| 142 | Frontal pole inferior | LimbicB | Limbic | 11 |
| 162 | Superior frontal gyrus mid-posterior lateral RH | DorsAttnB | Dorsal Attention | 11 |
| 185 | Middle frontal sulcus RH | ContB | Frontoparietal | 11 |
| 195 | Frontomarginal gyrus RH | ContB | Frontoparietal | 11 |
| 255 | Precuneus anterior LH | DefaultA | Default | 11 |
| 289 | Ventromedial prefrontal cortex posterior RH | DefaultA | Default | 11 |
| 295 | Central operculum RH | SalVentAttnA | Salience | 11 |
| 325 | Superior frontal gyrus superior RH | SalVentAttnA | Salience | 11 |
| 342 | Occipital pole inferior medial RH | VisCent | Visual | 11 |
| 344 | Middle temporal gyrus posterior RH | ContA | Dorsal Attention | 11 |
| 367 | Dorsomedial prefrontal cortex anterior RH | DefaultA | Default | 11 |
| 390 | Posterior cingulate cortex postero-inferior | DefaultA | Default | 11 |
| 421 | Middle temporal gyrus middle inferior LH | ContB | Default | 11 |
| 422 | Posterior orbital gyrus LH | SalVentAttnB | Frontoparietal | 11 |
| 423 | Paracingulate sulcus anterior LH | DefaultB | Default | 11 |
| 428 | Middle frontal gyrus anterior LH | SalVentAttnB | Frontoparietal | 11 |
| 442 | Cerebellum Crus II | No network found | No network found | 11 |
| 451 | Middle frontal gyrus middle inferior RH | ContB | Frontoparietal | 11 |

| | | | | |
|---|---|---|---|---|
| 467 | Callosomarginal sulcus superior LH | SomMotA | Somatomotor | 11 |
| 474 | Middle temporal gyrus middle RH | DefaultB | Default | 11 |
| 493 | Posterior cingulate cortex | DefaultA | Default | 11 |
| 495 | Intraparietal sulcus medial LH | ContA | Frontoparietal | 11 |
| 509 | Paracingulate sulcus middle LH | ContB | Frontoparietal | 11 |
| 523 | Precuneus middle superior RH | ContC | Dorsal Attention | 11 |
| 527 | Anterior insula superior RH | SalVentAttnB | Salience | 11 |
| 556 | Superior frontal sulcus anterior RH | DefaultA | Default | 11 |
| 572 | Paracingulate gyrus mid-posterior LH | SalVentAttnB | Frontoparietal | 11 |
| 633 | Middle frontal gyrus middle superior RH | ContB | Frontoparietal | 11 |
| 638 | Superior frontal gyrus mid-anterior LH | SalVentAttnB | Salience | 11 |
| 645 | Dorsomedial prefrontal cortex superior RH | ContB | Frontoparietal | 11 |
| 666 | Ventromedial prefrontal cortex | DefaultA | Default | 11 |
| 681 | Precuneus posterior LH | ContC | Frontoparietal | 11 |
| 683 | Anterior orbital gyrus RH | ContB | Frontoparietal | 11 |
| 704 | Anterior vertical ramus of the Lateral fissure RH | SalVentAttnB | Salience | 11 |
| 735 | Pars triangularis posterior LH | SalVentAttnA | Salience | 11 |
| 741 | Anterior cingulate cortex anterior LH | DefaultA | Default | 11 |
| 758 | Anterior cingulate cortex | SalVentAttnB | Frontoparietal | 11 |
| 788 | Middle frontal sulcus middle LH | SalVentAttnB | Salience | 11 |
| 796 | Inferior frontal gyrus middle LH | ContA | Frontoparietal | 11 |
| 824 | Subcentral gyrus anterior RH | SalVentAttnA | Salience | 11 |
| 843 | Callosomarginal sulcus mid-inferior | SalVentAttnA | Salience | 11 |
| 844 | Middle frontal sulcus inferior LH | ContB | Frontoparietal | 11 |
| 863 | Lateral occipital cortex mid-superior RH | DefaultC | Dorsal Attention | 11 |
| 870 | Precentral sulcus inferior LH | ContA | Dorsal Attention | 11 |
| 885 | Middle temporal gyrus posterior inferior LH | ContA | Dorsal Attention | 11 |
| 892 | Angular gyrus mid-posterior RH | DefaultA | Default | 11 |
| 916 | Pars opercularis inferior LH | SalVentAttnA | Salience | 11 |
| 947 | Callosal sulcus posterior | ContC | Frontoparietal | 11 |
| 980 | Inferior frontal gyrus anterior LH | ContB | Frontoparietal | 11 |

| | | | | |
|---|---|---|---|---|
| 991 | Ventromedial prefrontal cortex anterior LH | DefaultA | Default | 11 |
| 993 | Cerebellum Crus I LH | No network found | No network found | 11 |
| 1000 | Cingulate gyrus mid-anterior | SalVentAttnA | Salience | 11 |
| 6 | Putamen inferior RH | No network found | Subcortical | 12 |
| 24 | Cingulate sulcus mid-posterior LH | SalVentAttnA | Salience | 12 |
| 38 | Thalamus LH | No network found | Subcortical | 12 |
| 80 | Putamen posterior RH | SomMotB | Subcortical | 12 |
| 131 | Thalamus lateral LH | No network found | Subcortical | 12 |
| 172 | Parahippocampal gyrus posterior LH | No network found | Default | 12 |
| 183 | Postcentral sulcus inferior medial LH | DorsAttnB | Dorsal Attention | 12 |
| 184 | Anterior insula posterior RH | SalVentAttnA | Salience | 12 |
| 186 | Superior frontal gyrus posterior LH | DorsAttnB | Dorsal Attention | 12 |
| 251 | Caudate posterior | No network found | Subcortical | 12 |
| 262 | Midbrain superior | No network found | Subcortical | 12 |
| 281 | Thalamus middle | No network found | Subcortical | 12 |
| 290 | Cerebellum VI | No network found | No network found | 12 |
| 292 | Thalamus posterior RH | No network found | Subcortical | 12 |
| 385 | Cerebellum VI LH | No network found | No network found | 12 |
| 417 | Cerebellum V | No network found | No network found | 12 |
| 432 | Cingulate sulcus mid-anterior | SalVentAttnB | Frontoparietal | 12 |
| 435 | Cerebellum IX | No network found | No network found | 12 |
| 441 | Thalamus anterior RH | No network found | Subcortical | 12 |
| 444 | Globus pallidus posterior | SomMotB | Subcortical | 12 |
| 453 | Cerebellum IV | No network found | No network found | 12 |
| 553 | Paracentral sulcus LH | SomMotA | Somatomotor | 12 |

| | | | | |
|---|---|---|---|---|
| 575 | Middle frontal gyrus medial LH | SalVentAttnB | Frontoparietal | 12 |
| 594 | Planum temporale posterior LH | SomMotB | Somatomotor | 12 |
| 602 | Cingulate mid-anterior | SalVentAttnA | Salience | 12 |
| 606 | Putamen middle LH | SalVentAttnA | Salience | 12 |
| 626 | Parahippocampal gyrus | DefaultC | Default | 12 |
| 629 | Putamen and globus pallidus anterior LH | No network found | Subcortical | 12 |
| 632 | Hippocampus posterior RH | No network found | Subcortical | 12 |
| 647 | Cerebellum IV | No network found | No network found | 12 |
| 672 | Cerebellum IV | No network found | No network found | 12 |
| 679 | Insula superior LH | SalVentAttnA | Somatomotor | 12 |
| 698 | Thalamus antero-superior | No network found | Subcortical | 12 |
| 714 | Cerebellum V | No network found | No network found | 12 |
| 732 | Cerebellum V | No network found | No network found | 12 |
| 746 | Midbrain posterior | No network found | Subcortical | 12 |
| 857 | Cerebellum IV LH | VisPeri | No network found | 12 |
| 893 | Cerebellum V | No network found | No network found | 12 |
| 897 | Cerebellum V | No network found | No network found | 12 |
| 931 | Cerebellum VI anterior | No network found | No network found | 12 |
| 975 | Globus pallidus anterior | No network found | Subcortical | 12 |
| 982 | Cerebellum IX anterior | No network found | No network found | 12 |
| 1016 | Hypothalamus | No network found | Subcortical | 12 |
| 18 | Postcentral sulcus middle LH | DorsAttnB | Dorsal Attention | 13 |
| 21 | Lateral fissure posterior limb LH | SalVentAttnA | Salience | 13 |
| 132 | Precentral gyrus mid-inferior LH | SomMotB | Somatomotor | 13 |
| 163 | Precentral sulcus superior medial LH | DorsAttnB | Dorsal Attention | 13 |
| 176 | Superior temporal gyrus posterior medial LH | SomMotB | Somatomotor | 13 |

| | | | | |
|---|---|---|---|---|
| 315 | Postcentral sulcus inferior LH | DorsAttnB | Dorsal Attention | 13 |
| 328 | Postcentral gyrus inferior RH | DorsAttnB | Dorsal Attention | 13 |
| 331 | Superior parietal lobule anterior RH | DorsAttnA | Dorsal Attention | 13 |
| 348 | Precentral sulcus mid-inferior LH | ContA | Frontoparietal | 13 |
| 354 | Precentral gyrus middle anterior LH | DorsAttnB | Dorsal Attention | 13 |
| 430 | Postcentral gyrus mid-inferior LH | DorsAttnB | Dorsal Attention | 13 |
| 439 | Postcentral gyrus mid-superior RH | SomMotA | Somatomotor | 13 |
| 526 | Paracingulate gyrus posterior RH | SalVentAttnA | Salience | 13 |
| 542 | Postcentral sulcus middle RH | DorsAttnB | Dorsal Attention | 13 |
| 557 | Lateral fissure posterior limb inferior RH | SomMotB | Somatomotor | 13 |
| 585 | Superior frontal gyrus medial mid-posterior LH | SalVentAttnA | Salience | 13 |
| 589 | Lingual gyrus inferior LH | VisPeri | Visual | 13 |
| 609 | Central sulcus middle posterior RH | SomMotA | Somatomotor | 13 |
| 620 | Postcentral gyrus mid-superior LH | SomMotA | Somatomotor | 13 |
| 646 | Lateral occipital sulcus mid-anterior RH | DorsAttnA | Visual | 13 |
| 650 | Paracentral lobule posterior | SomMotA | Somatomotor | 13 |
| 663 | Postcentral sulcus inferior RH | DorsAttnB | Dorsal Attention | 13 |
| 701 | Postcentral sulcus mid-superior RH | DorsAttnB | Dorsal Attention | 13 |
| 718 | Superior parietal sulcus inferior RH | DorsAttnA | Dorsal Attention | 13 |
| 722 | Precentral sulcus superior LH | DorsAttnB | Somatomotor | 13 |
| 821 | Parietal operculum posterior RH | SalVentAttnA | Salience | 13 |
| 837 | Frontoparietal operculum LH | SomMotB | Somatomotor | 13 |
| 864 | Paracentral sulcus inferior | SomMotA | Somatomotor | 13 |
| 936 | Central sulcus middle RH | SomMotA | Somatomotor | 13 |
| 940 | Supramarginal gyrus anterior RH | DorsAttnB | Salience | 13 |
| 1011 | Superior parietal lobule inferior LH | DorsAttnB | Dorsal Attention | 13 |
| 461 | Precuneus postero-superior | ContC | Frontoparietal | 14 |
| 703 | Precuneus superior | ContC | Dorsal Attention | 14 |
| 748 | Precuneus interhemispheric fissure | ContC | Dorsal Attention | 14 |
| 941 | Parieto-occipital sulcus postero-superior | ContC | Visual | 14 |
| 55 | Middle temporal gyrus middle superior LH | DefaultB | Default | 15 |

| | | | | |
|---|---|---|---|---|
| 240 | Middle frontal gyrus posterior RH | ContB | Frontoparietal | 15 |
| 781 | Superior temporal gyrus posterior RH | TempPar | Somatomotor | 15 |
| 792 | Superior temporal gyrus middle LH | TempPar | Somatomotor | 15 |
| 925 | Superior temporal sulcus mid-posterior RH | TempPar | Somatomotor | 15 |
| 938 | Superior temporal gyrus mid-posterior RH | TempPar | Somatomotor | 15 |
| 10 | Calcarine cortex mid-posterior RH | VisPeri | Visual | 16 |
| 45 | Heschl's gyrus posterior RH | SomMotB | Somatomotor | 16 |
| 61 | Parieto-occipital sulcus posterior LH | ContC | Visual | 16 |
| 91 | Lateral occipital cortex antero-inferior RH | VisCent | Visual | 16 |
| 94 | Supracalcarine cortex RH | VisPeri | Visual | 16 |
| 102 | Occipital pole superior LH | VisCent | Visual | 16 |
| 137 | Cuneus posterior | VisCent | Visual | 16 |
| 155 | Occipital pole RH | VisCent | Visual | 16 |
| 158 | Calcarine cortex posterior LH | VisPeri | Visual | 16 |
| 247 | Fusiform gyrus mid-posterior LH | VisCent | Visual | 16 |
| 267 | Parieto-occipital sulcus posterior | VisPeri | Visual | 16 |
| 274 | Lateral occipital sulcus LH | DorsAttnA | Dorsal Attention | 16 |
| 323 | Calcarine cortex posterior RH | VisCent | Visual | 16 |
| 330 | Occipital pole inferior medial LH | VisCent | Visual | 16 |
| 343 | Lingual gyrus posterior inferior RH | VisCent | Visual | 16 |
| 350 | Occipital pole lateral LH | VisCent | Visual | 16 |
| 365 | Occipital pole LH | VisCent | Visual | 16 |
| 377 | Lunate sulcus inferior RH | VisCent | Visual | 16 |
| 380 | Occipital pole superior | VisCent | Visual | 16 |
| 383 | Occipital pole medial RH | VisCent | Visual | 16 |
| 427 | Parieto-occipital sulcus middle superior RH | ContC | Frontoparietal | 16 |
| 434 | Parieto-occipital sulcus mid-superior | ContC | Visual | 16 |
| 503 | Fusiform gyrus posterior LH | VisCent | Visual | 16 |
| 522 | Postcentral gyrus middle anterior LH | SomMotA | Somatomotor | 16 |
| 555 | Cuneus mid-posterior | VisPeri | Visual | 16 |
| 616 | Occipital pole | VisCent | Visual | 16 |

| | | | | |
|---|---|---|---|---|
| 665 | Lingual gyrus posterior LH | VisCent | Visual | 16 |
| 814 | Anterior occipital sulcus LH | VisCent | Visual | 16 |
| 839 | Retrocalcarine cortex LH | VisCent | Visual | 16 |
| 846 | Cuneus superior | VisPeri | Visual | 16 |
| 858 | Anterior occipital sulcus posterior RH | DorsAttnA | Visual | 16 |
| 865 | Precuneus postero-inferior LH | ContC | Frontoparietal | 16 |
| 899 | Occipital pole inferior | VisCent | Visual | 16 |
| 927 | Superior occipital gyrus mid-superior | VisPeri | Visual | 16 |
| 1007 | Cuneus superior RH | VisPeri | Visual | 16 |
| 1022 | Cuneus postero-inferior LH | VisPeri | Visual | 16 |
| 73 | Angular gyrus postero-inferior LH | TempPar | Default | 17 |
| 136 | Superior temporal sulcus mid-posterior LH | TempPar | Default | 17 |
| 144 | Superior temporal sulcus posterior RH | TempPar | Salience | 17 |
| 206 | Superior temporal sulcus middle RH | TempPar | Default | 17 |
| 238 | Lateral fissure anterior rami RH | SalVentAttnB | Salience | 17 |
| 242 | Supramarginal gyrus perisylvian LH | TempPar | Salience | 17 |
| 314 | Superior temporal sulcus posterior LH | TempPar | Somatomotor | 17 |
| 458 | Superior temporal sulcus middle LH | TempPar | Default | 17 |
| 545 | Angular sulcus anterior LH | TempPar | Salience | 17 |
| 596 | Angular sulcus mid-posterior LH | DefaultB | Default | 17 |
| 654 | Middle temporal gyrus mid-posterior RH | TempPar | Default | 17 |
| 769 | Superior temporal sulcus postero-superior RH | TempPar | Salience | 17 |
| 778 | Inferior frontal sulcus posterior RH | ContA | Frontoparietal | 17 |
| 828 | Superior temporal sulcus anterior LH | TempPar | Default | 17 |
| 852 | Angular gyrus antero-inferior LH | TempPar | Default | 17 |
| 972 | Middle temporal gyrus posterior superior LH | TempPar | Salience | 17 |
| 978 | Middle temporal gyrus mid-posterior LH | TempPar | Default | 17 |
| 1019 | Superior temporal gyrus anterior RH | TempPar | Somatomotor | 17 |
| 37 | Middle frontal gyrus posterior inferior RH | ContB | Frontoparietal | 18 |
| 118 | Inferior frontal gyrus posterior RH | ContA | Frontoparietal | 18 |
| 213 | Precentral sulcus middle RH | ContA | Dorsal Attention | 18 |

| | | | | |
|---|---|---|---|---|
| 214 | Angular sulcus anterior RH | TempPar | Salience | 18 |
| 218 | Pars triangularis RH | DefaultB | Frontoparietal | 18 |
| 249 | Angular sulcus mid-anterior LH | TempPar | Dorsal Attention | 18 |
| 285 | Superior temporal gyrus middle RH | SomMotB | Somatomotor | 18 |
| 311 | Superior temporal gyrus anterior | TempPar | Somatomotor | 18 |
| 318 | Superior frontal gyrus medial middle | SalVentAttnB | Salience | 18 |
| 351 | Superior occipital sulcus inferior LH | DorsAttnA | Dorsal Attention | 18 |
| 388 | Superior temporal sulcus mid-anterior LH | TempPar | Default | 18 |
| 408 | Inferior frontal sulcus posterior medial RH | ContA | Frontoparietal | 18 |
| 507 | Supramarginal gyrus postero-inferior RH | SalVentAttnA | Salience | 18 |
| 535 | Anterior thalamic radiation antero-superior LH | ContA | Frontoparietal | 18 |
| 674 | Pars opercularis lateral LH | DefaultB | Default | 18 |
| 752 | Occipital pole inferior LH | VisCent | Visual | 18 |
| 813 | Intraparietal sulcus inferior RH | ContA | Dorsal Attention | 18 |
| 818 | Inferior occipital sulcus inferior RH | VisCent | Visual | 18 |
| 854 | Supramarginal gyrus postero-inferior LH | SalVentAttnA | Salience | 18 |
| 888 | Precentral gyrus middle LH | DorsAttnB | Dorsal Attention | 18 |
| 963 | Precentral sulcus mid-superior LH | DefaultB | Frontoparietal | 18 |
| 1010 | Inferior frontal sulcus mid-posterior RH | ContA | Frontoparietal | 18 |
| 1018 | Superior temporal gyrus posterior lateral LH | TempPar | Salience | 18 |
| 31 | Lateral occipital cortex anterior RH | VisCent | Visual | 19 |
| 219 | Cuneus posterior RH | VisPeri | Visual | 19 |
| 284 | Occipital pole lateral RH | VisCent | Visual | 19 |
| 307 | Lateral occipital cortex inferior LH | VisCent | Visual | 19 |
| 373 | Lateral occipital cortex postero-superior LH | DorsAttnA | Visual | 19 |
| 468 | Lateral occipital cortex postero-inferior RH | VisCent | Visual | 19 |
| 695 | Superior occipital gyrus posterior LH | VisPeri | Visual | 19 |
| 719 | Cuneus postero-superior LH | VisPeri | Visual | 19 |
| 763 | Lateral occipital cortex middle RH | DorsAttnA | Dorsal Attention | 19 |
| 894 | Lateral occipital cortex middle LH | VisCent | Visual | 19 |
| 1013 | Lateral occipital sulcus anterior RH | DorsAttnA | Dorsal Attention | 19 |

| | | | | |
|---|---|---|---|---|
| 7 | Paracingulate gyrus posterior | SalVentAttnB | Salience | 20 |
| 32 | Middle frontal gyrus mid-posterior inferior RH | ContB | Frontoparietal | 20 |
| 66 | Middle frontal gyrus mid-anterior RH | SalVentAttnB | Frontoparietal | 20 |
| 85 | Cerebellum Crus I RH | No network found | No network found | 20 |
| 96 | Cingulate anterior | DefaultA | Default | 20 |
| 105 | Angular gyrus mid-superior RH | ContB | Default | 20 |
| 110 | Inferior frontal sulcus anterior LH | ContA | Frontoparietal | 20 |
| 120 | Cerebellum Crus I lateral LH | DorsAttnA | Dorsal Attention | 20 |
| 122 | Inferior occipital gyrus anterior LH | DorsAttnA | Dorsal Attention | 20 |
| 174 | Cerebellum Crus I superior RH | VisCent | Visual | 20 |
| 294 | Angular gyrus posterior superior RH | DefaultA | Default | 20 |
| 298 | Angular gyrus superior posterior LH | ContB | Default | 20 |
| 361 | Frontal pole inferior LH | ContB | Default | 20 |
| 366 | Ventrolateral prefrontal cortex RH | ContB | Frontoparietal | 20 |
| 369 | Frontal pole LH | LimbicB | Default | 20 |
| 405 | Paracingulate gyrus mid-anterior | DefaultA | Default | 20 |
| 419 | Intermediate primus of Jensen anterior RH | ContB | Frontoparietal | 20 |
| 463 | Pars orbitalis LH | DefaultB | Default | 20 |
| 470 | Intraparietal sulcus middle RH | ContA | Frontoparietal | 20 |
| 483 | Middle frontal gyrus middle superior LH | ContB | Default | 20 |
| 497 | Intermediate primus of Jensen LH | ContB | Frontoparietal | 20 |
| 506 | Superior frontal gyrus antero-superior RH | SalVentAttnB | Salience | 20 |
| 515 | Subparietal sulcus LH | DefaultA | Default | 20 |
| 548 | Superior frontal gyrus medial mid-anterior | ContB | Frontoparietal | 20 |
| 598 | Paracentral lobule postero-inferior RH | ContC | Salience | 20 |
| 600 | Gyrus rectus | LimbicB | Limbic | 20 |
| 652 | Superior frontal gyrus medial superior | DefaultB | Default | 20 |
| 700 | Supramarginal gyrus anterior LH | SalVentAttnA | Salience | 20 |
| 715 | Subparietal sulcus | DefaultA | Default | 20 |
| 730 | Middle frontal gyrus anterior RH | ContB | Frontoparietal | 20 |

| | | | | |
|---|---|---|---|---|
| 765 | Superior frontal gyrus anterior LH | DefaultB | Default | 20 |
| 787 | Dorsomedial prefrontal cortex antero-superior RH | DefaultB | Default | 20 |
| 802 | Superior frontal gyrus posterior lateral LH | DorsAttnB | Dorsal Attention | 20 |
| 831 | Middle frontal gyrus postero-superior LH | DefaultB | Default | 20 |
| 869 | Paracingulate gyrus middle | SalVentAttnB | Salience | 20 |
| 875 | Pars orbitalis RH | DefaultB | Default | 20 |
| 881 | Cerebellum Crus I RH | No network found | No network found | 20 |
| 903 | Precuneus anterior | ContC | Default | 20 |
| 918 | Subparietal sulcus posterior RH | ContC | Default | 20 |
| 929 | Middle frontal gyrus lateral LH | ContB | Frontoparietal | 20 |
| 944 | Frontal pole mid-superior RH | DefaultA | Default | 20 |
| 951 | Angular gyrus mid-anterior RH | SalVentAttnB | Default | 20 |
| 954 | Superior parietal lobule posterior | DorsAttnB | Dorsal Attention | 20 |
| 959 | Cerebellum Crus I posterior LH | VisCent | Visual | 20 |
| 984 | Angular gyrus anterior LH | DefaultB | Default | 20 |
| 994 | Middle frontal sulcus anterior LH | SalVentAttnB | Frontoparietal | 20 |
| 1015 | Superior frontal sulcus middle RH | ContB | Frontoparietal | 20 |
| 5 | Precentral sulcus mid-inferior RH | ContA | Dorsal Attention | 21 |
| 11 | Middle frontal gyrus superior RH | ContB | Default | 21 |
| 17 | Frontal pole lateral RH | ContB | Frontoparietal | 21 |
| 34 | Frontomarginal gyrus LH | DefaultA | Default | 21 |
| 126 | Ventrolateral prefrontal cortex LH | ContB | Frontoparietal | 21 |
| 129 | Supramarginal gyrus posterior RH | SalVentAttnB | Salience | 21 |
| 150 | Intraparietal sulcus anterior RH | ContA | Dorsal Attention | 21 |
| 188 | Angular gyrus postero-superior LH | DefaultC | Default | 21 |
| 224 | Superior occipital gyrus LH | DefaultC | Dorsal Attention | 21 |
| 253 | Supramarginal gyrus middle LH | SalVentAttnB | Salience | 21 |
| 254 | Middle frontal sulcus anterior RH | SalVentAttnB | Salience | 21 |
| 293 | Lateral orbital cortex LH | ContB | Frontoparietal | 21 |
| 336 | Precuneus superior RH | DorsAttnB | Dorsal Attention | 21 |

| | | | | |
|---|---|---|---|---|
| 353 | Intraparietal sulcus middle LH | ContA | Frontoparietal | 21 |
| 357 | Inferior temporal gyrus posterior LH | ContA | Frontoparietal | 21 |
| 374 | Supramarginal gyrus antero-superior RH | SalVentAttnA | Salience | 21 |
| 400 | Angular gyrus superior LH | DefaultA | Default | 21 |
| 412 | Superior frontal gyrus posterior lateral RH | ContB | Frontoparietal | 21 |
| 418 | Supramarginal gyrus superior RH | DorsAttnB | Salience | 21 |
| 437 | Middle temporal gyrus postero-inferior RH | DorsAttnB | Dorsal Attention | 21 |
| 443 | Paracingulate gyrus anterior | DefaultA | Default | 21 |
| 464 | Cerebellum Crus I LH | No network found | No network found | 21 |
| 492 | Frontal pole superior LH | DefaultB | Default | 21 |
| 494 | Pars triangularis inferior LH | DefaultB | Default | 21 |
| 496 | Angular gyrus antero-superior LH | DefaultB | Default | 21 |
| 514 | Inferior frontal sulcus anterior RH | SalVentAttnB | Frontoparietal | 21 |
| 525 | Intermediate primus of Jensen posterior RH | ContB | Frontoparietal | 21 |
| 528 | Angular gyrus middle RH | ContB | Frontoparietal | 21 |
| 532 | Inferior temporal gyrus middle RH | TempPar | Default | 21 |
| 533 | Superior rostral gyrus anterior | DefaultA | Default | 21 |
| 571 | Precuneus middle RH | ContC | Dorsal Attention | 21 |
| 642 | Frontal pole lateral LH | ContB | Frontoparietal | 21 |
| 656 | Frontal pole RH | LimbicB | Limbic | 21 |
| 667 | Ventromedial prefrontal cortex anterior | DefaultA | Default | 21 |
| 670 | Occipitotemporal sulcus mid-posterior RH | DorsAttnA | Visual | 21 |
| 728 | Pars triangularis anterior LH | ContA | Frontoparietal | 21 |
| 771 | Angular gyrus posterior LH | DefaultA | Frontoparietal | 21 |
| 776 | Middle frontal gyrus mid-anterior inferior RH | ContA | Frontoparietal | 21 |
| 806 | Middle frontal gyrus mid-anterior superior RH | SalVentAttnB | Salience | 21 |
| 809 | Lateral fissure posterior limb superior RH | SalVentAttnA | Salience | 21 |
| 880 | Ventromedial prefrontal cortex antero-inferior | LimbicB | Limbic | 21 |
| 884 | Supramarginal gyrus superior LH | ContA | Frontoparietal | 21 |
| 895 | Middle frontal gyrus lateral RH | SalVentAttnB | Frontoparietal | 21 |

| | | | | |
|---|---|---|---|---|
| 909 | Supramarginal gyrus posterior LH | DefaultB | Default | 21 |
| 934 | Frontal pole superior RH | DefaultB | Default | 21 |
| 945 | Supramarginal gyrus postero-superior LH | ContB | Frontoparietal | 21 |
| 955 | Callosomarginal sulcus mid-superior | DorsAttnB | Salience | 21 |
| 977 | Supramarginal gyrus posterior inferior RH | DefaultA | Default | 21 |
| 999 | Cerebellum Crus II RH | No network found | No network found | 21 |
| 1004 | Dorsomedial prefrontal cortex anterior LH | DefaultA | Default | 21 |
| 14 | Precentral gyrus middle | SomMotA | Somatomotor | 22 |
| 46 | Lingual gyrus posterior RH | VisPeri | Visual | 22 |
| 165 | Fusiform gyrus posterior RH | VisCent | Visual | 22 |
| 237 | Central sulcus inferior LH | SomMotB | Somatomotor | 22 |
| 268 | Postcentral gyrus mid-inferior lateral LH | SomMotA | Somatomotor | 22 |
| 381 | Postcentral gyrus inferior LH | SomMotB | Somatomotor | 22 |
| 510 | Central sulcus mid-inferior RH | SomMotB | Somatomotor | 22 |
| 539 | Heschl's gyrus anterior lateral RH | SomMotB | Somatomotor | 22 |
| 541 | Superior frontal gyrus medial posterior | SomMotA | Somatomotor | 22 |
| 621 | Lingual gyrus posterior | VisPeri | Visual | 22 |
| 636 | Postcentral gyrus middle RH | SomMotB | Somatomotor | 22 |
| 709 | Precentral gyrus inferior LH | SomMotB | Somatomotor | 22 |
| 868 | Middle parts of postcentral gyrus and precentral gyrus RH | SomMotA | Somatomotor | 22 |
| 956 | Central sulcus mid-inferior medial RH | SomMotB | Somatomotor | 22 |
| 971 | Central sulcus mid-inferior LH | SomMotB | Somatomotor | 22 |
| 997 | Subcentral gyrus posterior RH | SomMotB | Somatomotor | 22 |
| 13 | Superior parietal lobule anterior LH | DorsAttnA | Dorsal Attention | 23 |
| 121 | Fusiform gyrus mid-anterior RH | VisCent | Visual | 23 |
| 140 | Precentral sulcus superior RH | DorsAttnB | Dorsal Attention | 23 |
| 141 | Intraparietal sulcus anterior LH | ContA | Frontoparietal | 23 |
| 189 | Superior parietal lobule mid-posterior LH | DorsAttnA | Dorsal Attention | 23 |
| 387 | Superior occipital sulcus middle RH | DorsAttnA | Dorsal Attention | 23 |
| 448 | Superior parietal sulcus middle LH | DorsAttnA | Dorsal Attention | 23 |

| | | | | |
|---|---|---|---|---|
| 603 | Subcentral gyrus posterior LH | SomMotB | Salience | 23 |
| 611 | Lunate sulcus superior LH | VisCent | Visual | 23 |
| 686 | Intraparietal sulcus posterior LH | ContA | Dorsal Attention | 23 |
| 691 | Inferior temporal sulcus posterior LH | DorsAttnA | Dorsal Attention | 23 |
| 742 | Retrocalcarine sulcus LH | VisCent | Visual | 23 |
| 817 | Occipital pole inferior lateral RH | VisCent | Visual | 23 |
| 866 | Intraparietal sulcus mid-anterior LH | ContA | Frontoparietal | 23 |
| 878 | Superior parietal sulcus mid-inferior LH | DorsAttnA | Dorsal Attention | 23 |
| 905 | Lateral occipital sulcus mid-posterior RH | VisCent | Visual | 23 |
| 979 | Superior occipital sulcus superior RH | DorsAttnA | Visual | 23 |
| 995 | Superior occipital sulcus LH | DorsAttnA | Dorsal Attention | 23 |
| 998 | Superior parietal lobule posterior LH | DorsAttnA | Dorsal Attention | 23 |
| 20 | Parieto-occipital sulcus middle | DefaultC | Visual | 24 |
| 159 | Parieto-occipital sulcus middle RH | VisPeri | Visual | 24 |
| 445 | Parieto-occipital sulcus inferior LH | DefaultC | Visual | 24 |
| 583 | Posterior cingulate cortex inferior | DefaultC | Visual | 24 |
| 789 | Lingual gyrus mid-posterior LH | VisCent | Visual | 24 |
| 801 | Calcarine sulcus anterior | DefaultC | Default | 24 |
| 849 | Parieto-occipital sulcus | VisPeri | Visual | 24 |
| 939 | Parieto-occipital sulcus anterior LH | DefaultC | Default | 24 |
| 1020 | Parieto-occipital sulcus anterior RH | DefaultC | Visual | 24 |
